# Mapping the Morphology of DNA on Carbon Nanotube-Based Sensors in Solution using X-ray Scattering Interferometry

**DOI:** 10.1101/2023.05.04.539504

**Authors:** Daniel J. Rosenberg, Francis J. Cunningham, Joshua D. Hubbard, Natalie S. Goh, Jeffrey Wei-Ting Wang, Emily B. Hayman, Greg L. Hura, Markita P. Landry, Rebecca L. Pinals

## Abstract

Single-walled carbon nanotubes (SWCNTs) with adsorbed single-stranded DNA (ssDNA) are applied as sensors to investigate biological systems, with applications ranging from clinical diagnostics to agricultural biotechnology. Unique ssDNA sequences render SWCNTs selectively responsive to target analytes. However, it remains unclear how the ssDNA conformation on the SWCNT surface contributes to their ultimate functionality, as observations have been constrained to computational models or experiments under dehydrated states that differ substantially from the aqueous biological environments in which the nanosensors are applied. Herein, we demonstrate a direct mode of measuring in-solution ssDNA geometries on SWCNTs via X-ray scattering interferometry (XSI), which leverages the interference pattern produced by AuNP tags conjugated to ssDNA on the SWCNT surface. We employ XSI to quantify distinct surface-adsorbed morphologies for two ssDNA oligomer lengths, conformational changes as a function of ionic strength, and the mechanism of dopamine sensing for a previously established ssDNA-SWCNT nanosensor, with corresponding *ab initio* modeling for visualization. We show that the shorter oligomer, (GT)_6_, adopts a highly ordered structure of stacked rings along the SWCNT axis, compared to the longer, less periodic (GT)_15_ wrapping. The presence of dopamine elicits a simultaneous axial elongation and radial constriction of the ssDNA closer to the SWCNT surface. Application of XSI to probe solution-phase morphologies of nanoparticle-based tools will yield insights into sensing mechanisms and inform future design strategies for polymer-functionalized SWCNT technologies.

## Introduction

Single-walled carbon nanotubes (SWCNTs) serve as tools for biological sensing, imaging, and delivery applications. ^1,2^ SWCNTs are an advantageous platform due to their sensitive fluorescence response to localized changes (motivating sensor development^3–5^), photostable near-infrared fluorescence in the tissue-transparency window (enabling *in vivo* imaging^6,7^), and nanometer-sized diameter with a high aspect ratio (supporting use as cell-permeable delivery vehicles^8–10^). For each of these respective applications, the nanotube surface acts as a substrate upon which sensing moieties, anti-biofouling ligands, or delivery cargoes are loaded. Specifically, SWCNTs with adsorbed nucleic acids have been applied as nanoparticle-based sensors and delivery agents. Polymer properties including nucleic acid sequence and length govern SWCNT-adsorbed morphology, stability, and function. These constructs have proven particularly useful as nanosensors for small-molecule analytes including catecholamines^11–14^, serotonin,^15,16^ hydrogen peroxide,^17–19^ and nitric oxide^20,21^.

Despite over a decade of development in SWCNT-based sensors, there remain contrasting theories in the field about what enables molecular recognition, and what role (if any) conformational shifts play over chemical mechanisms. For example, a particular sequence of single-stranded DNA (ssDNA) – a repeating motif of guanine and thymine (GT) – has enabled highly sensitive and spatially resolved dopamine detection from single neurons and in neuronal tissue.^11–14^ Hypothesized interaction mechanisms between the dopamine and GT oligomer include dual hydrogen bonding between the two hydroxyl groups of dopamine and the phosphate backbone of the ssDNA,^12^ a redox reaction,^3^ and/or intercalation of the aromatic catecholamine ring between the ssDNA oligomer and SWCNT surface driven by *π-π* stacking.^3^ Optimizing interactions of nucleic acids with SWCNTs is key to the success of these biotechnologies, yet challenges remain in directly measuring, in real time, how ssDNA-SWCNT sensors behave.

Current methods for characterizing ssDNA-SWCNT conformations involve a dehydrated sample immobilized on a two-dimensional substrate, despite SWCNT-based biotechnologies mainly being applied in the aqueous solution state. Such techniques include transmission electron microscopy (TEM) to visualize ssDNA-SWCNT morphology^22,23^ and atomic force microscopy (AFM) to determine dimensions and packing of biomolecules on SWCNTs,^8,24–26^ which has been previously demonstrated to be limited by adsorption biases introduced during sample preparation.^27^ Other physical properties such as hydrodynamic dimensions can potentially be extracted from dynamic light scattering (DLS) measurements on SWCNTs done in the solution state, however, rigorous optical scattering methods have not been well-adapted for non-spherical, high-aspect-ratio particles such as SWCNTs and cannot resolve fine-grained surface features such as nanometer-scale polymer packing.

Small-angle X-ray scattering (SAXS) has shown promise in revealing the morphology of SWCNT-based systems in solution.^28,29^ We have previously reported the use of SAXS to determine the in-solution structure of ssDNA-suspended SWCNTs interacting with blood plasma proteins.^30^ However, SAXS is a contrast measurement technique relying on the scattering intensity of the analyte (proportional to the square of the electron density) being significantly higher than that of the solution. Thus, materials of relatively low electron density such as carbon-based SWCNTs and ssDNA must be at sufficiently high concentrations for the signal to be above background. Accordingly, characterizing ssDNA-SWCNTs via SAXS require the use of SWCNT concentrations that exceed those actually applied in biological systems (0.1-5 mg/L).^1,11,31^ These elevated concentrations can lead to artifacts such as inter-tube bundling of the ssDNA-SWCNTs,^23,28^ which must be minimized to fully elucidate the morphology of individual ssDNA-functionalized SWCNT sensors. A strategy to overcome this concentration issue for low-scattering materials is to increase the X-ray exposure time, but this risks creating chemical changes in solution that can affect the sample under study.^32^ An alternative approach is to increase the electron density of the sample directly using high contrast materials such as gold nanoparticles (AuNPs) and then apply X-ray scattering interferometry (XSI), originally described by Mathew-Fenn *et al*.^33,34^ XSI leverages the interference patterns generated upon X-ray scattering between ordered AuNPs to measure discrete inter-AuNP distances, effectively turning the AuNPs into molecular rulers in solution.^33–39^ Additionally, through adaptation of robotics and a rapid data-processing pipeline, XSI can be run at higher throughput with minimal sample consumption (∼1 sample per min with 30 μL per sample).^39–41^

Herein, we apply XSI to investigate nanopatterning of the adsorbed ssDNA corona surrounding the SWCNT surface in the solution phase. Small AuNPs (5.9-7.2 nm diameter) are attached to the 5’ end of each ssDNA oligomer^38,39^ and the ssDNA is adsorbed to the SWCNT surface, forming an ssDNA-SWCNT suspension with one AuNP tag per ssDNA strand. We focus on an illustrative example of how surface-constrained polymer conformation influences sensor properties by studying two (GT)_n_ ssDNA sequences (n = 6, 15) used for dopamine sensing. These two ssDNA oligomers empirically possess different advantageous properties, with (GT)_6_ displaying a larger magnitude of fluorescence change in response to dopamine^42^ and (GT)_15_ displaying higher stability in relevant biomolecule-rich environments.^43^ Previous characterization by both experimental studies (AFM, EM)^22,24^ and molecular dynamics (MD) simulations^42,44^ has led to postulation that these two oligomers possess distinct surface-constrained conformations: the shorter (GT)_6_ oligomer is expected to form a ring-like structure around the nanotube and the longer (GT)_15_ oligomer is expected to form a helical wrapping around the nanotube. We employ high-throughput XSI in solution at biologically applicable concentrations to explore: (i) the configuration of adsorbed ssDNA along the SWCNT surface, (ii) the conformational changes of adsorbed ssDNA as a function of ionic strength, and (iii) the behavior of adsorbed ssDNA in the presence of the target analyte, dopamine. Additionally, we perform *ab initio* modeling of the AuNPs adsorbed on the SWCNT surface directly from scattering profiles to provide a more comprehensive, 3D view of the system. We validate our technique with other suspension characterization (absorbance, fluorescence, DLS) and direct visualization (TEM). Taken together, this approach establishes a high-throughput technique for in-solution characterization of these nanoparticle-based biotechnologies and provides a deeper understanding surrounding the mechanisms behind their molecular recognition.

## Results and Discussion

### Synthesis and characterization of ssDNA-AuNP-SWCNTs

To first demonstrate the resolvable concentration range for (GT)_15_ ssDNA and (GT)_15_-SWCNTs without AuNP tags, we collected SAXS profiles of serial dilutions for each (Figure S1A-B). Bundled ssDNA-SWCNTs were observed even at the lowest resolvable concentration of 16 mg/L, with average bundling of ∼6-8 SWCNTs obtained from the cross-sectional radius of gyration (*R*_cs_; see SI Methods). No scattering contribution from the ssDNA alone was detected at equivalent concentrations (4 μM). These results and electron density calculations (see SI Methods and Extended Discussion) motivate our use of small AuNP tags (5.9-7.2 nm diameter) to increase the electron density of our material and thus observe ssDNA-SWCNTs in solution at relevant applied concentrations (<5 mg/L SWCNTs). Absolute-scale intensity scattering measurements demonstrate that scattering from 6.9 nm diameter PEGylated AuNPs (PEG-AuNPs) is ∼270-fold higher than that of (GT)_15_-SWCNTs and ∼65,000-fold higher than that of (GT)_15_ ssDNA (Figure S1C) at the correct relative concentrations (250 nM AuNP and ssDNA per 1 mg/L SWCNT; see SI Methods). Citrate-capped AuNPs were synthesized, conjugated to ssDNA via trithiolated linkers (Letsinger’s type) on the 5’ end, and coated with methoxy polyethylene glycol thiol (mPEG-SH) as detailed in SI Methods. Prepared ssDNA-AuNPs are then purified by anion exchange chromatography (Figure S2), characterized by SAXS to determine morphology and polydispersity (Figure S3 and Table S1), and DLS to confirm PEGylation by hydrodynamic radius (Figure S3 and Table S2) as detailed in SI Methods and Extended Discussion. As anticipated, the measured scattering in the conjugated systems of ssDNA-AuNP-SWCNTs is dominated by the AuNP signal and eliminates the need to mathematically factor in the scattering contributions from the ssDNA or SWCNT alone, or the scattering cross-terms between the different components of the complex.

We apply XSI to study two ssDNA sequences based on their relevance to biomolecular sensing and predicted surface-adsorbed conformational differences: (GT)_15_ and (GT)_6_.^11,12,22,42^ SWCNTs were suspended with ssDNA-AuNPs by probe-tip sonication at a constant ssDNA:SWCNT ratio (250 nmol ssDNA-AuNP per 1 mg SWCNT), in line with previous literature.^11,45^ We optimized this suspension method with the added AuNP tag, as detailed in the SI Methods and Extended Discussion. The resulting suspensions were characterized by absorbance and fluorescence analyses to corroborate ssDNA-AuNP-SWCNT complex formation (Figure S4). Retention of the AuNP plasmon resonance peak at approximately 520 nm reveals that the AuNP tags remain intact and well-dispersed through the SWCNT complexation process (Figure S4A). The apparent absence of absorbance peaks associated with SWCNT excitation is due to the high ratio of AuNPs to SWCNT in the ssDNA-AuNP-SWCNT complex and the limited dynamic range of the UV-Vis detector. Fluorescence spectra for AuNPs alone (with or without ssDNA) at 721 nm laser excitation reveal a trough in the emission intensity centered at approximately 950 nm. This optical feature may be due to absorption of excitation light, despite the lack of a distinct absorption band at this location. Compared to ssDNA-SWCNTs alone, addition of the AuNP tags results in lower intensity, broadened SWCNT fluorescence emission peaks, more prominently for (GT)_6_-than (GT)_15_-AuNP-SWCNTs and for emission peaks at shorter wavelengths (Figure S4B-D). This quenching effect of the AuNPs on the SWCNT fluorescence indicates electronic or excitonic interaction, and underscores the proximity of AuNPs to the SWCNT surface: AuNPs are metallic with known ultra-efficient quenching properties within 1-10s of nanometer-scale separation distances^46^ and SWCNTs are sensitive to perturbations in their local dielectric environment.^47^ The quenching mechanism may implicate photo-induced electron transfer or field effects of the proximal AuNPs on SWCNT excitons biasing toward nonradiative decay pathways,^48^ with potential vibrational contributions to peak broadening. The greater degree of quenching and peak-broadening observed for the (GT)_6_-AuNP-SWCNTs in comparison to the (GT)_15_-AuNP-SWCNTs likely arises from different ssDNA surface packing, where more (GT)_6_ ssDNA strands are expected per SWCNT based on previous literature.^30,49^ This difference between (GT)_15_- and (GT)_6_-AuNP surface packing is also confirmed by our TEM analysis as 0.139 vs. 0.185 AuNPs per nanometer length of SWCNT, respectively (see SI Methods). The relative enhancement of fluorescence at longer wavelength peaks suggests more large-diameter SWCNTS individually dispersed with the ssDNA-AuNPs.

### Conformational geometries of ssDNA on SWCNTs from XSI

The ssDNA-AuNP-SWCNT complexes were next analyzed by XSI. This technique is an extension of traditional solution SAXS in which a radial average of X-rays scattering off the electron density of a sample is integrated and the contrast between the sample and buffer is used to produce a buffer-subtracted 1D curve in reciprocal space. The total scattering intensity is the summation of two terms: the form factor, arising from the overall particle size and morphology, and the structure factor, derived from interparticle interactions. In structural biology, SAXS sample conditions (e.g., concentration) are adjusted to experimentally remove the contributions of the structure factor to isolate the form factor. An inverse Fourier transform of the subtracted curves then produces pairwise distribution functions, *P(r)*, providing real-space information on the average shape of the electron density of individual macromolecules free from interparticle interaction.^50,51^ Conversely, in XSI, the structure factor is of primary importance and can be isolated to represent the interference pattern of scattered X-rays arising from inter-AuNP interactions, indicating discrete distances between ordered AuNPs.^34,38,39^ In our case, we retain the form factor from the ssDNA-AuNP-SWCNT curves to enable normalization and additional analyses by preserving the information from the individual AuNPs.

Inter-ssDNA spacing along SWCNTs was measured using the AuNP tags via XSI (Figure 1A-B and SI Methods). Real-space analysis of the scattering profiles produces *P(r)* functions with two main peaks (Figure S5). The first peak represents the intra-AuNP distances between electrons within individual AuNPs, with the peak maximum being the average radius of the AuNPs. The absence of additional peaks in the *P(r)* functions without SWCNTs indicates that there is no long-range order and that the ssDNA-AuNPs (or PEG-AuNPs) are free in solution. The intra-AuNP peak provides a reference for the AuNP size distribution in each sample and enables normalization between samples to account for slight fluctuations in concentration and X-ray beam intensity. For clarity, the intra-AuNP peak is omitted in main figures but is included in supplementary figures. The second broader peak in the *P(r)* functions represents the inter-AuNP distances and is only observed in complexes containing periodic ordering of AuNPs. This peak reveals the distinct surface-adsorbed spacings of (GT)_15_- and (GT)_6_-AuNPs on the nanotube surface (Figure 1A-B). Importantly, ssDNA-AuNPs (without SWCNT substrates) are in a disordered state when free in solution and only enter a periodically ordered state when adsorbed to the SWCNT surface (Figure 1A-B, Figure S5, and Figure S6). TEM visualization recapitulates these findings in the dried state showing AuNPs ordered (Figure S7 and Figure S8) or free (Figure S9).

**Figure 1.**
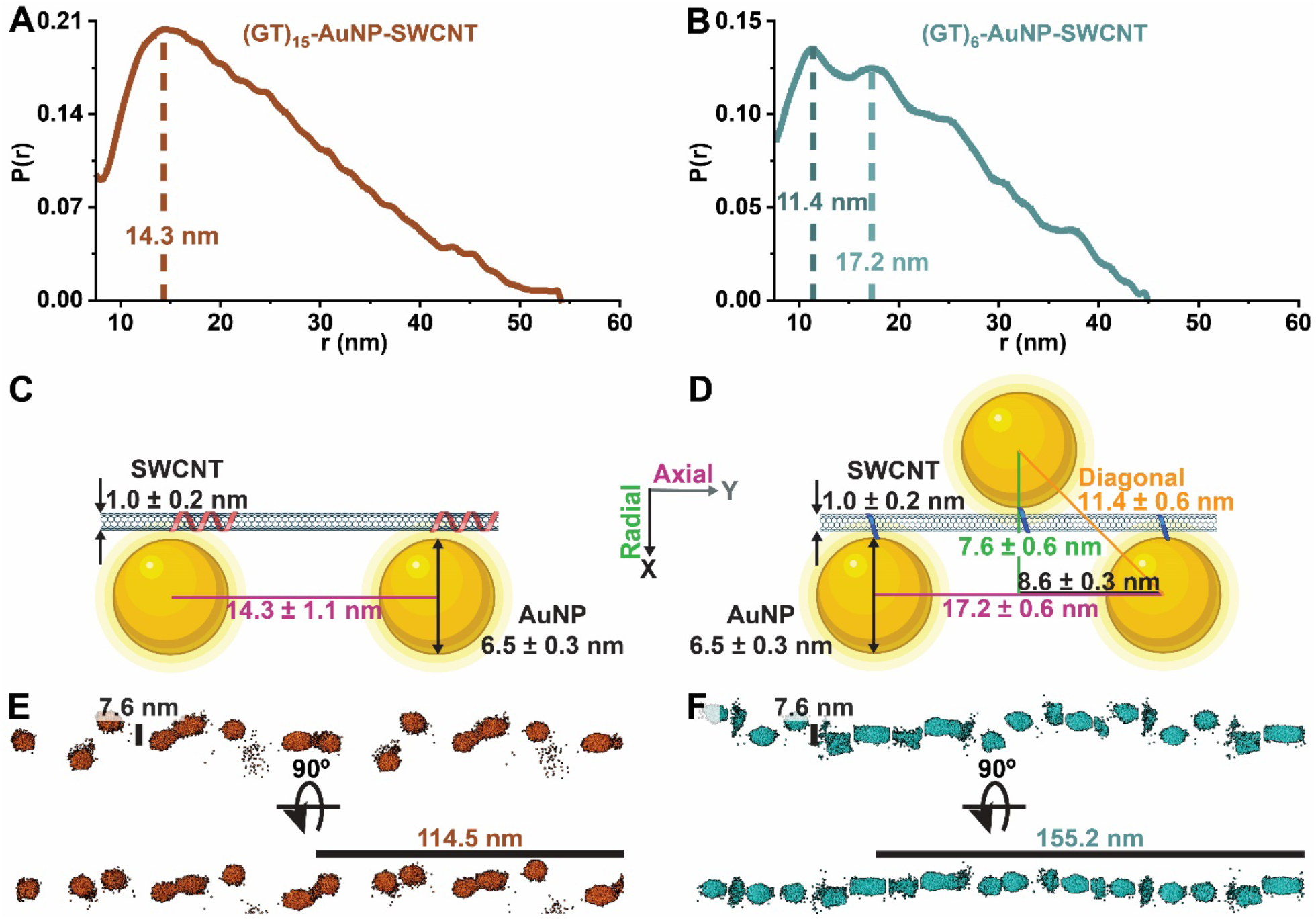
ssDNA forms ordered structures on the carbon nanotube surface. (A-B) Pairwise distribution functions, *P(r)*, from XSI data reveal discrete distances of AuNP-tagged ssDNA along the nanotube surface for (A) (GT)_15_-AuNP-SWCNTs and (B) (GT)_6_-AuNP-SWCNTs. *P(r)* functions are normalized to the primary intra-AuNP peak, then the x-axis minimum is set to focus on the inter-AuNP peak for clarity. (C-D) 2D schematics for proposed geometrical arrangement of AuNPs on the SWCNT surface, with average inter-AuNP distances obtained from statistical analysis of *P(r)* functions for (C) (GT)_15_-AuNP-SWCNT and (D) (GT)_6_-AuNP-SWCNT. Average inter-AuNP distances are denoted as diagonal (orange), axial (magenta), and radial (green). Schematics are drawn to scale. (E-F) *Ab initio* modeling results for (E) (GT)_15_-AuNP-SWCNT and (F) (GT)_6_-AuNP-SWCNTs. Fits and residuals are shown in Figure S13.

A series of controls was analyzed to confirm that the preparation of ssDNA-AuNP-SWCNT complexes leads to adsorption of ssDNA-AuNPs on the SWCNT surface rather than off-target aggregative process: ssDNA-SWCNTs (no AuNPs), free ssDNA-AuNPs (no SWCNTs), and carboxylated SWCNTs mixed with ssDNA-AuNPs (no probe-tip sonication, and thus no driving force for self-assembly) do not reveal any feature suggesting AuNP order (Figure S10A-B). Likewise, no order is observed in SWCNTs attempted-to-be suspended with PEG-AuNPs (no ssDNA) (Figure S10C-D). Additionally, there is no ssDNA-AuNP-SWCNT concentration dependence over the range used in this study (0.17-1.76 mg/L; Figure S11), which is within the unbundled SWCNT regime based on previous literature.^23^ Finally, there is no AuNP size dependence for the axial inter-AuNP distances over the range used in this study (5.9-7.2 nm diameter; Figure S12A-C). Therefore, at this ssDNA:SWCNT ratio (250 nmol ssDNA-AuNP per 1 mg SWCNT), the packing of ssDNA on the SWCNT surface is not affected by potential steric effects from the AuNPs.

MD simulations have shown that (GT)_15_ forms a helical wrapping around SWCNTs with uniform electrostatic potential profiles, as opposed to the ring-like conformation of (GT)_6_ showing a periodic electrostatic footprint.^42^ For (GT)_15_, this larger-footprint morphology and lower packing density (ascertained by TEM image analysis) is likely responsible for the generally broader and less defined inter-AuNP peaks observed for (GT)_15_-AuNP-SWCNTs and suggests a more variable surface adsorption pattern (Figure 1A and Figure S7). Due to this increased variability in (GT)_15_-AuNP adsorption and lack of orientational reference, a statistical analysis of the most probable inter-AuNP distances (14.3 ± 1.1 nm) is used to determine the basic 1D axial ssDNA spacing along the SWCNT (Figure 1C, Figure S12A, and C). In contrast, the inter-AuNP peaks for (GT)_6_-AuNP-SWCNTs are narrower and contain more clearly defined higher-order features after the initial, most probable distance (11.4 ± 0.6 nm; Figure 1B and Figure S12B-C). MD simulations of (GT)_6_ on (9,4) chirality SWCNTs (0.92 nm diameter) predict that there is a near-equivalent split in energetically favorable left-handed helix and ring-like conformations for low packing densities, and that steric effects from moderate-to-high surface coverage result in a population shift to primarily rings.^42^ Accordingly, (GT)_6_ is expected to adopt a ring-like configuration on the majority of SWCNTs used in this study (mixed chiralities, with average diameter of 1 ± 0.2 nm).

From the longer-range distance features of the inter-AuNP peak of (GT)_6_-AuNP-SWCNTs, we deduce that the AuNPs align on alternating sides of the SWCNT. The second-most probable inter-AuNP distance reveals an average center-to-center distance of 17.2 ± 0.6 nm (Figure 1B and Figure S12B-C), representing the inter-AuNP spacing axially down the SWCNT and establishing the preceding peak at 11.4 nm the distance of AuNPs diagonally across the SWCNT (Figure 1D). These inter-AuNP distances are used to extrapolate the average periodic inter-ssDNA ring distance of 8.6 ± 0.3 nm (Figure 1D). As expected, the axial inter-AuNP distances for both (GT)_15_- and (GT)_6_-AuNP-SWCNTs are not affected by changes in AuNP diameter (Figure S12C). Conversely, the diagonal distance is expected to change slightly as a function of the AuNP diameter, as calculated in Figure S12C when the AuNPs are flush with the SWCNT surface, holding to a 2D geometry (deemed adequate due to the large diameter disparity between AuNPs and SWCNTs). This trend is not seen experimentally, however, suggesting that the AuNPs are not directly in contact with the SWCNT surface, and their positional variance may preclude our ability to see this trend. To explore these dynamics, diagonal and axial inter-AuNP distances from a single size of AuNPs (d = 6.1 ± 0.03 nm) were used to calculate an average radial distance of 7.6 ± 0.6 nm (Figure S12D). This means that, using this simplified 2D geometry, the average distance from SWCNT surface to nanoparticle surface is 0.8 ± 0.3 nm.

We expand upon these 2D geometric analyses with 3D visualization of the ssDNA-AuNPs adsorbed on the SWCNT surface through *ab initio* modeling directly from scattering profiles using SASHEL (see SI Methods and Extended Discussion). The final best-fit models demonstrate the complexity of these systems and validate our 2D interpretation (Figure 1E-F and Figure S13A-C). As expected, *ab initio* models for (GT)_6_-AuNP-SWCNTs show increased AuNP packing density and greater consistency in inter-AuNP distances than (GT)_15_-AuNP-SWCNTs. When rotated about the SWCNT axis, (GT)_6_-AuNP-SWCNTs reveal a plane where there is little radial variance between AuNPs (Figure 1F), confirming our hypothesis that the small diameter of the SWCNT would reasonably support a 2D estimate of geometries. Conversely, no such plane was found for (GT)_15_-AuNP-SWCNT models and the AuNPs seem to rotate freely around the SWCNT axis (Figure 1E).

Our calculated separation distances of ssDNA polymers along the SWCNT axis are in agreement with previous literature.^52^ Based on prior MD simulations, a single (GT)_15_ polymer footprint on a (9,4) chirality SWCNT is expected to extend ∼4 nm in length, with a ∼2 nm helical pitch.^42^ Another MD simulation-based study similarly estimates the pitch of (GT)_30_ oligonucleotides on (11,0) SWCNTs to be 2-8 nm, depending on the DNA backbone orientation (with the 8 nm pitch orientation more energetically favorable, albeit on larger diameter SWCNTs).^53^ This latter modeling study determined that a previously measured 18 nm pitch helix of poly-(GT) strands around SWCNTs via atomic force microscopy (AFM)^54^ was structurally unstable and most likely introduced as an artifact during the air-drying step necessary for AFM sample preparation. Another AFM-based study suggests ∼14 nm pitch for (GT)_15_ on SWCNTs.^27^ DNA pitch on SWCNTs has also been visualized by TEM, with estimates of 2.2 nm pitch for double-stranded salmon testes DNA along SWCNTs.^55^ These previous pitch estimates can be converted to inter-strand spacing as measured in the current study via geometrical calculation, with an average SWCNT diameter of 1 nm and 0.7 nm length per ssDNA base,^24,27^ assuming that inter- and intra-strand pitch distances are equivalent. Additionally, adjacent ssDNA strands are assumed to be close, but not intertwined, along the SWCNT axis.^24^ Pitch estimates from the aforementioned literature ranging from 2-18 nm correspond to (GT)_15_ inter-strand spacing of 15.7-20.9 nm and (GT)_6_ inter-strand spacing of 6.3-8.4 nm. The directly measured inter-strand values in solution of 14.3 ± 1.1 nm and 8.6 ± 0.3 nm for (GT)_15_- and (GT)_6_-AuNP-SWCNT, respectively, are reasonable in comparison to those computationally predicted or measured in previous studies.^52,56^

### Surface-adsorbed ssDNA structural changes as a function of ionic strength

We applied this XSI approach to determine *in situ* ssDNA packing on the SWCNT surface as a function of solution ionic strength (Figure 2). Increasing solution ionic strength is expected to modify the surface-adsorbed ssDNA conformation, and thus AuNP scattering periodicity, by screening the negatively charged phosphate backbone of the ssDNA and enabling closer packing along the nanotube surface.^57,58^ To test this hypothesis, ssDNA-AuNP-SWCNTs were synthesized, dialyzed against 0.1X PBS, and then diluted to various PBS concentrations to achieve different net salt concentrations while maintaining constant pH (see SI Methods). We tested samples in a range of 0.05X to 2X PBS represented as corresponding Debye lengths (λ_D_) ranging from 3.37 to 0.53 nm, calculated as previously described.^59^ This range was selected because ssDNA-SWCNTs are less stable in pure water and PBS concentrations above 2X resulted in aggregation.

**Figure 2.**
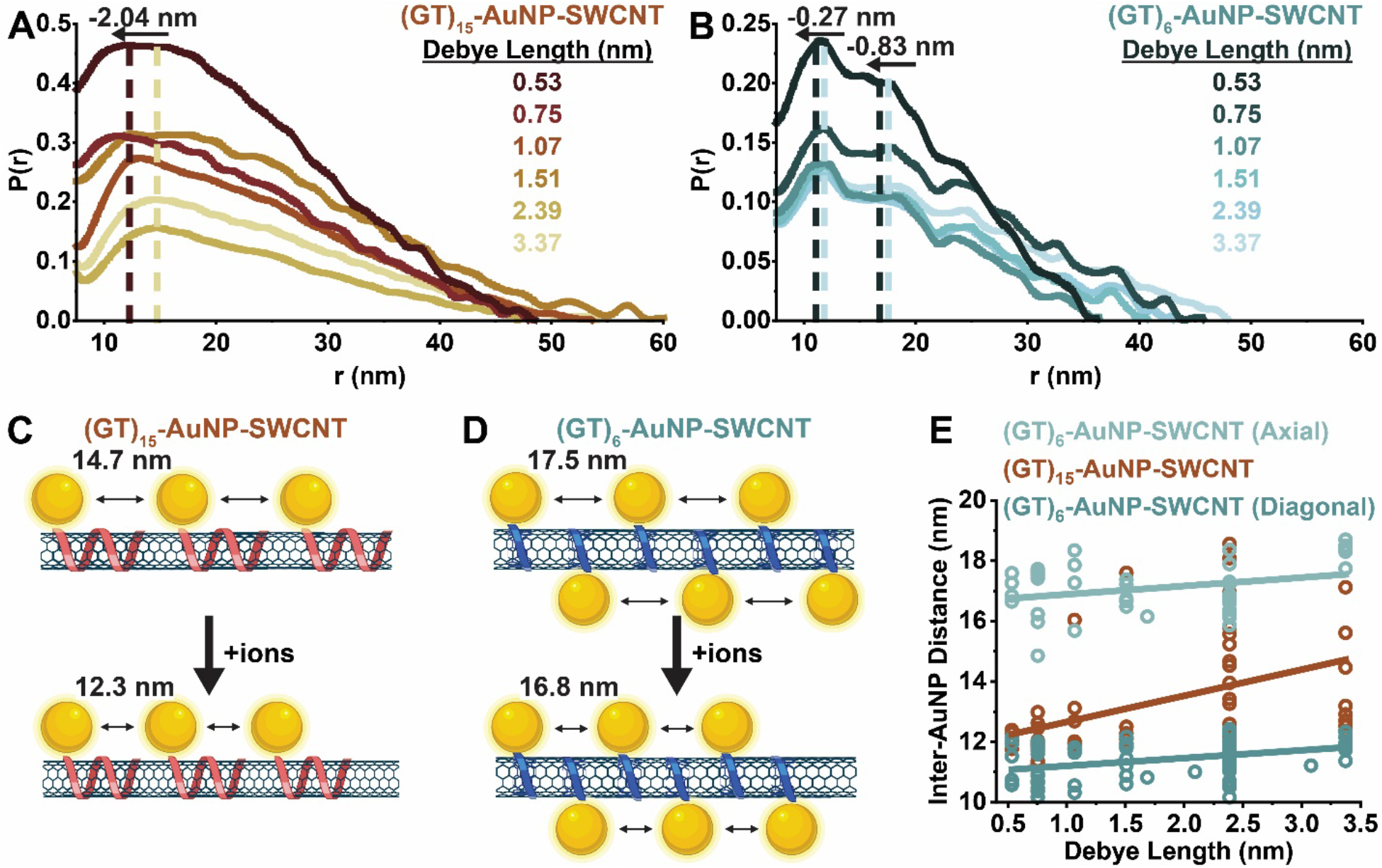
Surface adsorbed inter-ssDNA distance is modulated as a function of ionic strength for longer polymer lengths but remains relatively unchanged for shorter polymer lengths. Representative pairwise distribution functions, *P(r)*, for (A) (GT)_15_-AuNP-SWCNTs (red-orange series) and (B) (GT)_6_-AuNP-SWCNTs (blue series) in phosphate-buffered saline of varying net salt concentration, as represented by Debye lengths (λ_D_ = 3.37-0.53 nm). Dashed vertical lines are added to visualize peak shifts proceeding from light to dark dashed lines. *P(r)* functions are normalized to the primary intra-AuNP peak, then the x-axis minimum is set to focus on the inter-AuNP peak for clarity. (C-D) Schematic representations of changes in inter-AuNP distances at elevated ion concentrations for (C) (GT)_15_-AuNP-SWCNTs and (D) (GT)_6_-AuNP-SWCNTs. Schematics are not drawn to scale. Additional, representative *P(r)* functions and scattering curves are included in Figure S5 and Figure S6 for (GT)_15_- and (GT)_6_-AuNP-SWCNTs, respectively. (E) Summary of inter-AuNP distances as a function of Debye length (λ_D_ = 3.37-0.53 nm) for individual samples (dots) with corresponding linear regression (lines).

As predicted, the longer, multi-pass helices of (GT)_15_ on the nanotube surface compress at increased salt conditions (lower λ_D_) from inter-AuNP distances of 14.7 to 12.3 nm (Figure 2A, C, and E). Conversely, the spacing of the shorter, single-pass rings of (GT)_6_ did not change as significantly with ionic strength in either diagonal or axial inter-AuNP distances (Figure 2B, D, and E). An axial inter-AuNP shift from 17.5 to 16.8 nm was measured, corresponding to inter-ssDNA distances of 8.8 to 8.4 nm (for λ_D_ = 3.37-0.53 nm). Conversely, the calculated radial distances show negligible change as a function of ionic concentration (for λ_D_ = 3.37-0.53 nm; Figure S12D).

Given the smaller decrease in inter-AuNP distance observed for (GT)_6_-AuNP-SWCNT at increased ionic strengths compared to that of (GT)_15_-AuNP-SWCNT, we postulate that the high-salt condition affects the local intra-strand pitch to a greater extent than the neighboring inter-strand interactions. Moreover, increasing the concentration of ions in solution does not alter the radial distances of (GT)_6_-AuNPs across the SWCNT, as expected due to the short-range nature of the *π-π* interactions between the ssDNA and SWCNTs. Our salt-dependent ssDNA surface-packing results for (GT)_15_ spacing on SWCNTs are in line with previous literature demonstrating this phenomenon with longer ssDNA on SWCNTs via indirect optical measurement and dried-state characterization.^57,58^ At high salt concentrations, (GT)_30_ was determined to adopt a compact conformation with higher SWCNT surface coverage,^57^ putatively due to self-stacking of nucleobases from a related MD study.^60^ In comparison, the ssDNA enters an elongated and stiffer conformation at low salt concentration, accompanied by ssDNA desorption from the SWCNT reducing the packing density.^57^

### ssDNA-SWCNT nanosensor interactions with dopamine

We employed XSI to explore the ssDNA conformational changes of (GT)_15_- and (GT)_6_-AuNP-SWCNT complexes in the presence of the nanosensor target analyte, dopamine (DA) (Figure 3). Upon injection of 100 μM DA, a shift in the average inter-AuNP distances was observed in the *P(r)* functions for both ssDNA lengths. Inter-(GT)_15_ strand spacing increased by 2.11 ± 1.0 nm (Figure 3A, C, and Figure S14A), while the axial inter-(GT)_6_ strand spacing increased by only 0.59 ± 0.27 nm (as calculated from the axial inter-AuNP peak shift of 1.17 ± 0.55 nm) and the diagonal inter-AuNP peak revealed an average shift of -0.93 ± 0.11 nm (Figure 3B-C and Figure S14B). Based on this observation, we calculated a corresponding radial inter-AuNP distance shift of -2.2 ± 0.41 nm for (GT)_6_-AuNP-SWCNTs in the presence of DA, reducing the average SWCNT-to-AuNP-surface distance to -0.07 ± 0.04 nm. This dramatic decrease in radial distance demonstrates that DA causes the ssDNA to constrict around the SWCNT, drawing in the AuNP tags. Moreover, this slightly negative SWCNT-to-AuNP distance suggests that the AuNPs begin to overlap in the plane of the SWCNT and thus the ssDNA rings may be preferentially wrapping in opposite directions. Of note, this shortening of the radial inter-AuNP distances is not observed as a function of ionic strength (Figure S12D) and underscores the analyte-specific binding capabilities of this surface-constrained ssDNA sequence. *Ab initio* modeling of the AuNPs adsorbed on the SWCNT surface confirm the decrease in the average radial inter-AuNP distances for (GT)_6_-AuNP-SWCNTs in the presence of DA (Figure 3D and Figure S15).

**Figure 3.**
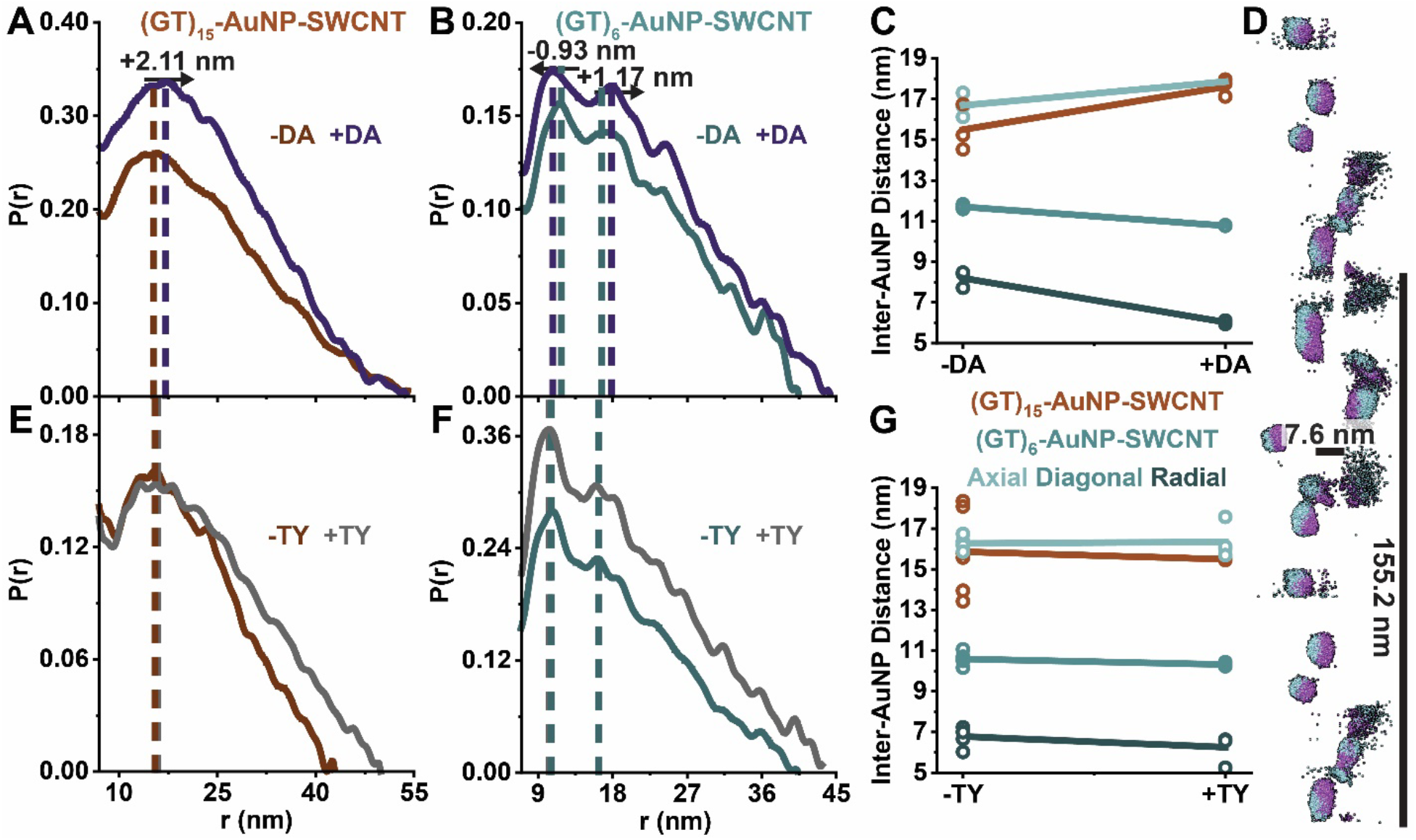
Inter-ssDNA distances shift in opposite directions for axial and radial spacing in the presence of dopamine and vary based on the ssDNA length and conformation. Inter-AuNP spacings shift in the presence of (A-D) dopamine (DA) but not with (E-G) *p*-tyramine (TY), a structural analog. Representative pairwise distribution functions, *P(r)*, with no analyte for (A and E) (GT)_15_-AuNP-SWCNTs (red-orange series) and (B and F) (GT)_6_-AuNP-SWCNTs (blue series) or in the presence of (A-B) DA (purple series) or (E-F) TY (grey series). Dashed vertical lines are added to visualize peak shifts. *P(r)* functions are normalized to the primary intra-AuNP peak, then the x-axis minimum is set to focus on the inter-AuNP peak for clarity. (C and G) Summary of inter-AuNP distances for replicates with and without (C) dopamine (DA) and (G) *p*-tyramine (TY). For (GT)_6_-AuNP-SWCNT samples, axial, diagonal, and radial inter-AuNP distances are shown for individual samples (dots) with corresponding linear regression (lines). Corresponding *P(r)* functions for replicates are shown in Figure S14. (D) *Ab initio* modeling results for (GT)_6_-AuNP-SWCNTs demonstrate the decrease in radial inter-AuNP distances as they move from blue to purple locations in the presence of DA. Fit and residuals are shown in Figure S15.

From the axial expansion of (GT)_n_ down the length of the SWCNT and additional radial constriction of (GT)_6_ onto the SWCNT, we postulate that DA both interacts with the phosphate groups of the ssDNA and inserts between ssDNA bases, depending on the initial conformation of the adsorbed polymers. Previous MD simulations of (8,8) chirality (GT)_15_-SWCNTs (d = 1.1 nm) in the presence of DA suggest that the hydroxyl groups of DA (protonated at pH 7.4) interact with the exposed phosphate groups of the ssDNA backbone, drawing the ssDNA closer to the SWCNT surface.^12^ Another MD study of (9,4) chirality (GT)_15_- and (GT)_6_-SWCNTs (d = 0.92 nm) reported that DA inserted between bases of the helically wrapped (GT)_15_, but failed to insert between the bases of the ring-like (GT)_6_, forming bridges between neighboring strands instead.^42^ Both proposed mechanisms show that the DA interaction creates localized perturbations in the periodically ordered, ssDNA-induced electrostatic surface potentials of the SWCNT. These perturbations modulate exciton recombination lifetimes and lead to a large increase in nanosensor fluorescence. Our observation that the axial inter-strand distances of (GT)_15_-AuNP-SWCNTs increase to a greater extent than (GT)_6_-AuNP-SWCNTs complements the hypothesis that DA preferentially inserts between the bases for (GT)_15_, increasing the pitch and hence footprint length along the SWCNT surface. Interestingly, as observed in (GT)_6_-AuNP-SWCNTs, the presence of DA also constricts the ring-like structure around the SWCNT, suggesting that DA interaction is pulling the phosphate backbone of ssDNA towards the SWCNT surface. This interaction may also be the case for (GT)_15_-SWCNTs, but no radial distances can be calculated due to the lack of orientational reference. As a control, XSI was collected for ssDNA-AuNP-SWCNTs in the presence of the dopamine analogue, *p*-tyramine (TY), containing only one hydroxyl group. Negligible changes in the inter-ssDNA distances were observed upon injection of TY (Figure 3E-G and Figure S14C-D), consistent with the lack of fluorescence response and predictions from MD simulations.^11,42^

## Conclusions

In this work, we demonstrate that XSI is a valuable technique for studying (GT)_n_-SWCNTs (n = 6, 15) in solution, using small AuNP tags conjugated to the ssDNA to act as molecular rulers. XSI harnesses the tightly packed, electron-rich gold atoms in AuNPs to enable the study of nanomaterials at concentrations relevant to biological applications (0.1-5 mg/L). We find periodic ordering of ssDNA-AuNPs along the SWCNT axis, with the highest probability inter-AuNP distance of 14.3 ± 1.1 nm for (GT)_15_-AuNPs and 11.4 ± 0.6 nm for (GT)_6_-AuNPs. For (GT)_6_-AuNP-SWCNTs, higher-order features observed after the most probable inter-AuNP distance motivated more detailed geometric calculations, giving rise to an extrapolated average inter-ssDNA ring distance of 8.6 ± 0.3 nm and an average distance from SWCNT-to-AuNP surface of 0.8 ± 0.3 nm.

Exploration of *in situ* ssDNA packing on the SWCNT surface as a function of solution ionic strength over the range of 0.05X to 2X PBS (λ_D_ = 3.37-0.53 nm) reveals an inter-ssDNA spacing decrease for (GT)_15_-AuNP from 14.7 to 12.3 nm and (GT)_6_-AuNP from 8.8 to 8.4 nm. These observations reflect the predicted electrostatic charge screening of the ssDNA phosphate backbone to permit closer packing. From these findings we postulate that the high-salt condition has a greater effect on the local intra-strand pitch rather than on the neighboring inter-strand interactions. The minimal change in radial AuNP spacing as a function of solution ionic strength suggests a lesser role of electrostatics in driving ssDNA-SWCNT adsorptive interactions, as expected for the likely *π*-*π* and/or other hydrophobic forces governing the polymer-surface adsorption mechanism.

XSI elucidates the conformational changes of (GT)_15_ and (GT)_6_ ssDNA adsorbed on SWCNTs in the presence of the nanosensor target analyte, DA, and provides insight into the mechanism responsible for the increased fluorescence response. Prior to analyte addition, radiative recombination of excitons is quenched by closely packed (GT)_6_ rings that create periodic positive and negative surface potential pockets on the SWCNT.^42,61^ Our observations demonstrate a perturbation of these predicted surface potentials in the presence of dopamine, as the axial distance is expanded between the strands and the ssDNA is pushed closer to the SWCNT surface, leading to an increase in radiative recombination pathways and/or decrease in nonradiative decay mechanisms. In the presence of DA, inter-ssDNA spacing increases by 2.11 ± 1.0 nm for (GT)_15_-AuNPs and by only 0.59 ± 0.27 nm for (GT)_6_-AuNPs. The greater shift in inter-ssDNA for (GT)_15_-AuNP indicates that DA inserted between bases occur to a more significant degree for the helically wrapped (GT)_15_, increasing the pitch and hence the footprint length on the SWCNT surface as previously predicited.^42^ Interestingly, for (GT)_6_-AuNP-SWCNTs the radial inter-AuNP distances show a dramatic decrease of 2.2 ± 0.41 nm in the presence of DA, which is not detected in response to changes in salt concentrations. This change is confirmed by *ab initio* modeling and demonstrates a constricting of the (GT)_6_ ring likely caused by the hydroxyl groups of DA interacting with the exposed phosphate groups of the ssDNA, pulling the ssDNA closer to the SWCNT surface. This constriction leads to a calculated SWCNT-to-AuNP surface distance of -0.07 ± 0.04 nm, implying that AuNPs overlap in the plane on one side of the SWCNT. This potential geometry suggests that the ssDNA rings have coordinated directionality, wrapping in opposite directions from each other along the SWCNT surface. These findings attest to the complexity of these nanobiotechnologies, suggesting that the mechanisms behind their molecular recognition of DA is conformationally driven, and hence sequence-specific, leading to a strong argument towards the need for rational design to properly tailor their optical properties.

As demonstrated, XSI provides a powerful tool to complement previously employed techniques to characterize DNA-based nanotechnologies under biologically relevant solution-phase conditions. Through this technique, we gain understanding of the discrete nanoscale architectures of these materials and their mechanisms of interaction with the local environment in solution. Such high-throughput measurements are important to understand how polymer-nanoparticle complexes function for a broad range of nanobiotechnology applications.

## Acknowledgements

J.D.H. acknowledges the support of the National Science Foundation. N.S.G. acknowledges financial support from the Foundation for Food and Agriculture Research (FFAR) Fellows program. We acknowledge support of the IGI LGR ERA, GlaxoSmithKline, and Citris/Banatao Seed Funding. We acknowledge support of a Burroughs Wellcome Fund Career Award at the Scientific Interface (CASI) (to M.P.L.), a Dreyfus foundation award (to M.P.L.), an NIH MIRA award (to M.P.L.), an NSF CAREER award (to M.P.L), an NSF CGEM award (to M.P.L.), a FFAR Young Investigator award (to M.P.L.), a CZI investigator award (to M.P.L), a Sloan Foundation Award (to M.P.L.), a and Moore Foundation Award (to M.P.L.). M.P.L. is a Chan Zuckerberg Biohub investigator, a Hellen Wills Neuroscience Institute Investigator, and an IGI Investigator. R.L.P. acknowledges support from the Schmidt Science Fellows program, in partnership with the Rhodes Trust. Efforts to apply XSI for studying ssDNA on SWCNTs are supported in part by National Cancer Institute grants Structural Biology of DNA Repair (SBDR) CA092584 and CA220430. XSI data was collected at the Advanced Light Source (ALS) beamline SIBYLS which is supported by the DOE-BER IDAT DE-AC02-05CH11231 and NIGMS ALS-ENABLE (P30 GM124169 and S10OD018483). This work benefited from the use of the SasView application, originally developed under NSF award DMR-0520547. SasView contains code developed with funding from the European Union’s Horizon 2020 research and innovation program under the SINE2020 project, grant agreement No 654000. We thank the staff at the University of California, Berkeley Electron Microscope Laboratory for advice and assistance in electron microscopy sample preparation and data collection. Additional electron microscopy was conducted at the National Center for Electron Microscopy at the Molecular Foundry, Lawrence Berkeley National Laboratory. Work at the Molecular Foundry was supported by the Office of Science, Office of Basic Energy Sciences, of the U.S. Department of Energy under Contract No. DE-AC02-05CH11231. Molecular graphics and analyses performed with UCSF Chimera, developed by the Resource for Biocomputing, Visualization, and Informatics at the University of California, San Francisco, with support from NIH P41-GM103311. We would like to acknowledge the use of medical clipart from BioRender.com. Thank you to Dr. Daniel Murray, Dr. Lee Joon Kim, Dr. Abraham Beyene, Dr. James Holton, Dr. Andrew Crothers, and Dr. Michal Hammel for their helpful feedback and editing of this manuscript. Thank you to Brandon Russel for enabling remote modeling operations. Thank you to Elizabeth Voke for making this long-distance project possible by always being ready to send images of written lab notebooks during the manuscript writing process.

## Author contributions

Conceptualization, D.J.R. and R.L.P.; methodology (XSI, SAXS), D.J.R.; investigation (XSI, SAXS), D.J.R.; investigation (TEM), J.D.H. and D.J.R.; investigation (DLS), D.J.R. and N.S.G.; investigation (absorbance, fluorescence), N.S.G. and R.L.P.; formal analysis (XSI, SAXS), D.J.R.; formal analysis (TEM), D.J.R.; formal analysis (DLS), D.J.R. and N.S.G.; formal analysis (absorbance, fluorescence), N.S.G. and R.L.P.; writing, D.J.R. and R.L.P.; sample preparation, D.J.R., R.L.P., N.S.G., J.W.W., and F.J.C.; coding, D.J.R., E.B.H. and R.L.P.; supervision, M.P.L., G.L.H., and R.L.P.; project administration, M.P.L. and G.L.H.; funding acquisition, M.P.L. and G.L.H.

