## Supplementary Information for "Mapping the Morphology of DNA on Carbon Nanotube-Based Sensors in Solution using X-ray Scattering Interferometry"

### Methods

#### Materials

5'-trithiolated-ssDNA (Letsinger's type) oligonucleotides with SDS-PAGE purification were purchased from Fidelity Systems (Gaithersburg, MD). Methoxy poly(ethylene glycol) thiol (mPEG-SH; MW ~350 g/mol) was purchased from Biochempeg Scientific Inc. (Watertown, MA). Small diameter raw HiPco™ single-walled carbon nanotubes (SWCNTs) were purchased from NanoIntegris (Boisbriand, Quebec, Canada). Carboxylated SWCNTs were purchased from Sigma Aldrich (Burlington, MA). All other reagents were purchased from Millipore Sigma (St. Louis, MO).

#### Synthesis of Citrate-Capped Gold Nanoparticles (AuNPs)

Citrate-capped AuNPs of diameters 5.9-7.2 nm were prepared using a method modified from that which was previously described.[1] Briefly, a 2 L solution of 0.25 mM $HAuCl_4$ and 0.25 mM tri-sodium citrate was prepared in a conical flask using dd$H_2O$ cooled to 4°C. Next, 10 mL of 0.6 M $NaBH_4$ at 4°C was added rapidly to the solution while stirring. The solution turned dark red immediately after adding $NaBH_4$, indicating particle formation. The solution was allowed to warm to room temperature (RT) and stirred overnight for the water to decompose excess $NaBH_4$.

#### Citrate-BSPP Exchange for Gold Nanoparticles (BSPP-AuNPs)

Bis-(*p*-sulfonatophenyl) phenylphosphine (BSPP) was added to citrate-stabilized colloidal AuNPs (~5.0 x $10^{13}$ particles/mL) to a final concentration of 0.5 g/L and stirred at RT for a minimum of 6 hours. Approximately 1 mL of saturated NaCl solution was added per 10 mL BSPP-exchanged colloidal gold, until the solution changed from transparent red to a darker, cloudy purple, indicating the reversible precipitation of the AuNPs.[2] The mixture was centrifuged (Beckman, JA-18 rotor) at 12,000 rcf for 10 min and decanted carefully to not disturb the pelleted BSPP-AuNPs. BSPP-AuNPs were washed twice with 0.5 M NaCl solution (repeating the centrifugation step above) and resuspended in 15 mM phosphate buffer, 1 mM TCEP, pH 7 for storage. Suspensions were stored at 4°C until use. Note that freezing caused sample precipitation.

#### Conjugation of Single-Stranded DNA to BSPP-AuNPs (ssDNA-AuNPs)

BSPP-AuNPs were attached to ssDNA via trithiolated linkers (Letsinger's type) on the 5' end of ssDNA oligomers, then coated with short, neutral methoxy polyethylene glycol thiol (mPEG-SH) polymers and purified, as previously described.[3,4] If stored longer than 2 weeks, fresh TCEP was added to reduce the solution of colloidal BSPP-AuNPs prior to ssDNA conjugation. To do this, saturated NaCl solution was first added to BSPP-AuNPs until the solution turned dark, then the BSPP-AuNPs were centrifuged at 12,000 rcf for 10 min and resuspended in 15 mM phosphate buffer, 1 mM TCEP, pH 7. The final BSPP-

AuNP concentration was determined by measuring the absorbance at 520 nm (NanoDrop 2000, Thermo Scientific) and converting to concentration with the empirical extinction coefficient,[5] $\varepsilon_{520nm}$ = 9.69x$10^6$ L $mol^{-1}$ $cm^{-1}$. The concentration of desired 5'-trithiolated-ssDNA (Letsinger's type) oligonucleotides was calculated by measuring the absorbance at 260 nm using the sequence-dependent extinction coefficient. Solutions of BSPP-AuNPs and trithiolated-ssDNA were mixed vigorously at a final molar ratio of 1:1 and incubated at RT overnight. A 129 mM mPEG-SH solution was prepared and added to the ssDNA-AuNP suspension at a final molar ratio of 3000:1 mPEG-SH to AuNPs. The final ssDNA-AuNP concentration was determined again by measuring absorbance.

#### Anion Exchange Chromatography Purification of ssDNA-AuNPs

Mono-conjugated ssDNA-AuNPs were isolated as previously described[3] using a Dionex DNA-Pac PA100 anion exchange column on either a GE AKTA Explorer or a GE Atka Pure fast protein liquid chromatography (FPLC) with an NaCl gradient from 0.01 to 1 M over a period of 55 min at a flow rate of 1.1 mL/min. Sample elution was monitored by measuring UV-Vis absorption at 260 (ssDNA) and 520 nm (AuNPs), and the mono-conjugated ssDNA-AuNP fraction was collected for downstream use (Figure S2A). The same AuNP conjugation and purification method was implemented for both $(GT)_{15}$ and $(GT)_6$ oligomers, with the longer demonstrating increased retention times via anion exchange chromatography, as expected (Figure S2B).

#### Characterization by High-Throughput X-ray Scattering Interferometry (HT-XSI)

HT-XSI data was collected at the SIBYLS beamline (bl12.3.1) at the Advanced Light Source of Lawrence Berkeley National Laboratory, Berkeley, California.[6] X-ray wavelength was set at $\lambda$ = 0.12398 nm and the sample-to-detector distance was 2.07 m, resulting in a scattering vector ($q$) range of 0.1 - 4.6 $nm^{-1}$, which corresponds to real-space distances of 62.8 - 1.4 nm. The scattering vector is defined as $q = 4\pi\sin\theta/\lambda$, with scattering angle $2\theta$. Data was collected using a Dectris PILATUS3X 2M detector at 20°C and processed as described previously.[7]

Immediately prior to data collection, 15 µL of each sample was added to 15 µL of buffer in a 96-well plate kept at 10°C for final corresponding concentrations of 42.5 - 450 nM ssDNA-AuNPs and 0.17 - 1.76 mg/L SWCNTs. Each sample was then transferred to the XSI sampling position via a Tecan Evo liquid handling robot (Tecan Trading AG, Switzerland) with modified pipetting needles acting as sample cells as described previously.[8] Samples were exposed to X-ray synchrotron radiation for 5 s at a 0.1 s frame rate for a total of 50 images. Each collected image was circularly integrated and normalized for beam intensity to generate a one-dimensional scattering profile by beamline-specific software (Blu-Ice). Buffer subtraction was performed for the one-dimensional scattering profile of each sample using each of two bracketing buffer wells to ensure the subtraction process was not subject to instrument variations. Scattering profiles over the 5 s exposure were sequentially averaged together to eliminate any potential radiation damage effects. Averaging was performed by batch processing using the HT-XSI data processing pipeline in the SIBYLS SAXS Process (SSP) GUI.[4,9] Pairwise distribution functions, *P(r)*, were generated in batch using the automated GNOM[10] feature of the SSP GUI. Average SWCNT bundling of ~6-8 SWCNTs determined by calculating the cross-sectional radius of gyration ($R_{cs}$) for rod-like scatterers using ATSAS 3.0.[11] Absolute-scale intensity scattering measurements and calculations were completed as previously described.[12]

All AuNPs used in this study are near-spherical, with average diameters ranging from 5.9 to 7.2 nm determined from pairwise distribution functions (*P(r)*; probability plot of all inter-electron distances) obtained from SAXS for each batch of synthesized ssDNA-AuNPs (Figure S3A, Table S1, and Table S2). SAXS profiles for all ssDNA-AuNPs were further modeled as triaxial ellipsoidal fittings[13] using SasView software to obtain more accurate nanoparticle dimensions (Figure S3B, Table S1, Table S2, and detailed in SI Extended Discussion). The average ratio of largest (major equatorial radius, $r_A$) to smallest (polar radius, $r_C$) particle dimension is observed at 3.3:2. Including a polydispersity parameter for $r_A$ was essential to optimize the fit (Figure S3C). The average polydispersity index (PDI) of $r_A$ is 0.18, indicating reasonable monodispersity, where PDI < 0.1 is considered ideal.[14]

#### Electron Density Calculations

For X-ray scattering experiments, the electron density of a material is of critical importance, as the total scattering intensity is proportional to the square of the electron density. The triaxial ellipsoidal AuNPs used in this study have a calculated electron density of 3519.08 $e^-/nm^3$, increasing the electron density of our otherwise low-scattering materials with calculated electron densities of 914.3 and 1818 $e^-/nm^3$ for SWCNTs and ssDNA, respectively. Detailed electron density calculation are found in SI Extended Discussion.

#### Characterization by Dynamic Light Scattering (DLS)

DLS measurements were taken with the Zetasizer Nano ZS (Malvern Analytical) with a material refractive index of 0.200 and absorption of 3.320 for colloidal gold.[15,16] All samples were diluted in 0.1X PBS to an AuNP concentration of 0.20-0.25 μM and loaded in disposable cuvettes (Malvern ZEN0040) for size measurement.

AuNPs were coated with mPEG-SH to prevent aggregation and aid in the purification by anion exchange chromatography.[3] After mPEG-SH coating, the hydrodynamic radii of the AuNPs measured by DLS increase from the AuNP core radii measured with SAXS by an average of 1.46 ± 0.34 nm across all samples (Figure S3D and Table S3). This increase in hydrodynamic diameter implies successful mPEG-SH functionalization of the exposed AuNP surfaces.[17] The average length of the mPEG-SH (MW ~350 g/mol) is ~1.7 nm when fully extended, based on the length of the repeating polyethylene oxide unit of 0.278 nm in water (with n=6 units total).[18] However, even if all available mPEG-SH functionalized the surface, the theoretical distance between grafting sites (~5.2 nm) would far exceed the calculated Flory radius of 0.815 nm, leading to only a partially extended mushroom conformation. This result is in line with the measured change in hydrodynamic radius imparted by mPEG-SH being slightly less than the expected change based on polymer length.[19–21]

#### Suspension of Single-Walled Carbon Nanotubes (SWCNTs) with ssDNA-AuNPs

Purified mono-conjugated ssDNA-AuNPs were spin-concentrated prior to use (Amicon Ultra-0.5 mL centrifugal filters with 3 kDa MWCO, Millipore Sigma): 500 µL of ssDNA-AuNP solution was centrifuged in the filter at 14,000 rcf for 20 min, the filtrate was removed to waste, and an additional 450 µL of ssDNA-AuNP solution was added, repeating the centrifugation at 14,000 rcf for 20 min. ssDNA-AuNP sample (approximately 10X concentrated) was recovered by reversing the filter into a clean tube and centrifuging at 1,000 rcf for 2 min. ssDNA-AuNP concentration was determined by measuring the absorbance at 520 nm (NanoDrop One, Thermo Scientific) with a 10X-diluted aliquot and calculating the concentration as before.

Single-walled carbon nanotube (SWCNTs) were suspended with ssDNA-AuNPs as follows: mixed-chirality raw SWCNTs (small diameter HiPco™ SWCNTs, raw, NanoIntegris) were first prepared as an aqueous slurry of 2 mg/mL in Milli-Q water. ssDNA-AuNP-SWCNTs were then formulated to maintain a final ratio of 250 nmol ssDNA-AuNP per 1 mg SWCNT, at a total volume of 2-4 mL (such that half of the solution could serve as non-sonication controls). The exact formulation recipe depended on the yield of ssDNA-AuNPs obtained after anion exchange purification. For every 2 mL of ssDNA-AuNPs at 200-800 nM (0.4-1.6 nmol), 0.8-3.2 µL of SWCNT slurry was added (1.6-6.4 µg) in 0.1X phosphate-buffered saline (PBS; note 1X PBS is 137 mM NaCl, 2.7 mM KCl, 10 mM $Na_2HPO_4$, 1.8 mM $KH_2PO_4$, pH 7.4) in a 5 mL tube. The ssDNA-AuNP/SWCNT mixture was bath sonicated for 10 min (Branson Ultrasonic 1800) then probe-tip sonicated for 10 min in an ice bath (3 mm probe tip at 50% amplitude, 5-6 W, Cole-Parmer Ultrasonic Processor). ssDNA-AuNP-SWCNT suspension was equilibrated for 30 min at RT then dialyzed against 2 L of 0.1X PBS overnight (200 µL volume in Pur-A-Lyzer Mini Dialysis Kit with 6-8 kDa MWCO, Millipore Sigma). Note that free ssDNA-AuNPs are not expected to pass through this filter size that contains pores of only a few nanometers and this step was included as a buffer exchange to remove any impurities still present from the AuNP synthesis. Suspensions were stored at 4°C until use. Control experiments without SWCNTs were prepared from the same batch of ssDNA-AuNPs, with all steps the same but in the absence of SWCNTs.

Certain parameters were slightly modified in comparison to usual ssDNA-SWCNT suspension protocols to account for the AuNP tag on the ssDNA. First, the material amounts were reduced approximately two orders of magnitude to account for the limited availability of AuNPs, although kept in a similar ratio to previous suspension protocols (250 nmol ssDNA:1 mg SWCNT).[22,23] Second, a post-sonication pelleting step, which one would typically perform in order to remove unsuspended SWCNTs or amorphous carbon and catalyst left over from SWCNT synthesis, was omitted because the presence of the AuNPs causes full sample pelleting due to the additional mass. We expect that reducing the overall material load by two orders of magnitude also reduced the concentration of SWCNT-derived impurities in the final suspension, thus eliminating the need for a centrifugation clean-up step. Finally, SWCNT suspensions were not spin-filtered due to embedding of the AuNPs into the filter membrane and full sample loss. Additional suspension notes are detailed in the SI Extended Discussion.

#### Characterization by Absorbance and Fluorescence

Absorbance was measured with a UV-VIS-nIR spectrophotometer (Shimadzu UV-3600 Plus) using 50 µL sample volume in black-sided quartz cuvette (Thorlabs, Inc.). Near-infrared SWCNT fluorescence was measured using an inverted Zeiss microscope (Axio Observer.D1, 10x objective) with a Princeton Instruments spectrometer (SCT 320) and liquid nitrogen-cooled Princeton Instruments InGaAs detector (PyLoN-IR). A triggered 721 nm laser (OptoEngine LLC) was used as the excitation source and fluorescence emission was collected from 800 – 1400 nm. 30 µL volume of each sample was prepared in polypropylene 384 well-plates (Greiner Bio-One microplate).

#### Characterization by Transmission Electron Microscopy (TEM)

Images of $(GT)_6$-AuNP-SWCNT and $(GT)_{15}$-AuNP-SWCNT complexes were captured using a Tecnai 12 TEM (FEI, Hillsboro, OR) operating at an accelerating voltage of 120 kV and data was recorded using a Gatan Rio16 CMOS camera with GWS software (Gatan Inc., Pleasanton, CA). Samples were prepared by depositing 5 µL of sample onto 400 mesh carbon/formvar-coated copper grids (EMS Electron Microscopy Science) that were surface treated by glow discharge to render the support hydrophilic. The samples were wicked away after 2 minutes. No negative staining or washing steps were included. Approximately 20 images were taken at four different regions on the grid for each sample to ensure reported images were representative. Additional image analysis was performed using Fiji (Imagej2).[24] To determine the size distribution and average AuNP size, over

1000 AuNPs were analyzed. To calculate the AuNP packing on the SWCNTs, a minimum of 5 SWCNTs were analyzed per sample giving a total SWCNT length of 2980 ± 62 nm.

#### Modeling SAXS Data

Modeling of the SAXS profiles was done using SASHEL, adapting the methodology originally described by Burian and Amenitsch.[25] SASHEL was first developed as an algorithm to reconstruct helical and rod-like systems by randomly moving dummy atoms comprising a single building block unit and projecting them outward using symmetrical boundary conditions. Theoretical scattering profiles from these projected models are iteratively fit to experimental data until convergence on the best-fit model. In a typical SASHEL analysis, it is recommended to reduce the number of total SAXS datapoints uniformly across the original 1D curve (i.e., removing every-other datapoint) to lower calculation times. However, this leads to a higher density of datapoints in the high q-range due to the binning of the circular integration of the detector images and reduces the fitting quality in the low q-range. In this study, datapoints were sequentially removed at higher density as the q-value increases to retain high fit quality in the low q-range, corresponding to the longer order distances of interest. The data was fit over a truncated q-range of 0.1-3 $nm^{-1}$ to further focus on the lower q-range.

Initial conditions for the model were refined over many iterations, with the most robust outcome derived from a carefully curated starting model for both $(GT)_{15}$- and $(GT)_6$-AuNP-SWCNT samples (Figure S16). First, the more clearly defined $(GT)_6$-AuNP-SWCNT scattering curve was used to produce a crude initial estimate of dummy-atom positions (Figure S16A). This model started with a core-shell cylinder model with 15 nm outer diameter and 5 nm inner diameter, as determined by the average diameter of the SWCNT (~1 nm) together with a fully extended trithiolated linker on either side (~2 nm each). The stack building block height ($H_{BB}$) was set equal to the axial inter-AuNP distance (17.2 nm) of $(GT)_6$-AuNP-SWCNT, as determined from *P(r)* functions (Figure 1B). The number of stacks ($N_S$) was set to 15 to ensure an overall length far greater than the observed maximum length dimension, $d_{max}$, of the $(GT)_6$-AuNP-SWCNT samples (~55 nm). The starting temperature ($T_0$) was set to 0.6 to allow for broad movement of the 2000 initial dummy atoms per $H_{BB}$. $T_0$ in this case is the value in which the system starts to cool down as defined previously[25] and is not representative of temperature on an absolute scale. Dummy-atom diameters were set to 0.288 nm to simulate the atomic diameter of a gold atom. From this initial model, clusters of dummy atoms formed and were taken to be naturally representative of AuNPs (regions of high electron density). The most clearly defined cluster was extracted, duplicated to create a pair of AuNPs, and the number of dummy atoms expanded to 1000 for each AuNP (Figure S16A).

Each AuNP in the initial pair is maneuvered into estimated initial geometries as determined experimentally to represent the inter-AuNP block heights ($H_{GB}$) unique for $(GT)_{15}$- and $(GT)_6$-AuNP-SWCNTs (14.3 and 17.2 nm, respectively). Each refined AuNP pair ($H_{GB}$ unit) was then replicated axially using symmetrical boundary conditions an integer ($k$) number of times to produce a total stack height, $H_{BB} = k*H_{GB}$, as depicted in Figure S16B and C for $(GT)_{15}$- and $(GT)_6$-AuNP-SWCNTs, respectively. After exploring a large parameter space, (outlined in detail in SI Extended Discussion, Table S4, Table S5, Figure S17, Figure S18, Figure S19, and Figure S20) it was discovered that when so few dummy atoms were used per AuNP, the models would fit better by expanding outward to compensate for a lack of representative electron density. Thus, initially modeling $(GT)_{15}$- and $(GT)_6$-AuNP-SWCNTs as two neighboring ssDNA-AuNP-SWCNTs saved a lot of computational time (less dummy atoms to move per iteration) and allowed for a larger parameter space to be explored. These inital models were run at a starting temperature of 0.2 at each of the respective $H_{BB}$ values, with $N_S$ = 32 / number of AuNPs ($N_{NP}$) rounded to the nearest integer value, where $N_{NP}$ is the total number of AuNPs per stack. All models were run for 200 iterations, as the goodness of fit ($\chi^2$-value) tends to reach a minimum plateau.

To model the system as a single SWCNT, the best fit double SWCNT model for $(GT)_{15}$- and $(GT)_6$-AuNP-SWCNT was selected (Figure S21) and the single SWCNT region showing the clearest AuNP spacing is extracted. The regions of clear electron density are replaced with denser 3000 dummy-atom clusters (representing AuNPs; Figure S16B-C) and an initial $H_{BB}$ was used which resulted in the best fit from each respective double SWCNT model ($H_{BB}$ = 114.5 and 155.2 nm for $(GT)_{15}$- and $(GT)_6$-AuNP-SWCNT respectively; Table S5 and Figure S20). The $N_S$ were 3 and 2 for $(GT)_{15}$- and $(GT)_6$-AuNP-SWCNT models, respectively, to obtain final models of similar total length (342.6 and 310.4 nm, respectively). Over many iterations, as regions in the model appeared which suggested areas of excessive or missing electron density, AuNPs (dummy-atom clusters) were manually removed or added accordingly until achieving a reasonably fitting single SWCNT model ($\chi^2$-value < 2; Figure S13A-B). Higher starting temperatures ($T_0$ > 0.4) were generally used for single SWCNT to speed up the movement of initial models containing at least three-fold more dummy atoms than earlier models.

### Extended Discussion

#### Electron Density Calculations

SWCNTs

The surface structure of a SWCNT is that of a sheet of graphene.[26] Assuming the C-C bond length $d_{c\text{-}c}$ = 0.1421 nm is the same for graphene as for a curved SWCNT, the surface area of one carbon hexagon ($a_h$) is:

$$a_h = \frac{3\sqrt{3}}{2} d_{c-c}^2 = 0.0525\, nm^2 \tag{1}$$

The calculated surface area of one benzene ring corresponds to two carbon atoms and thus the electron surface density of the hexagon ($\sigma$) is:

$$\sigma = \frac{\text{number electrons } (e^-)}{a_h} = 228.57\, e^-/nm^2 \tag{2}$$

The electron density ($\rho$) of a SWCNT is calculated as follows with surface area ($a_s$) = $\pi dL$ and volume ($V$) = $\pi(d/2)^2L$ for a SWCNT of average diameter ($d$) = 1 nm (according to technical data sheet from NanoIntegris) and arbitrary length ($L$):

$$\rho = \frac{a_s\sigma}{V} = \frac{4\sigma}{d} = 914.3\, e^-/nm^3 \tag{3}$$

AuNPs

The number of gold atoms is based on calculations from ellipsoidal AuNPs, as previously described.[27] The equation for calculating the number of atoms in a triaxial ellipsoidal AuNP ($N_{Au}$) is as follows:

$$N_{Au} = \frac{4\pi}{3V_{Au}}(r_A - d_{Au})(r_B - d_{Au})(r_C - d_{Au}) \tag{4}$$

Where $V_{Au}$ = 0.017 nm$^3$ is the volume of the gold atom and $d_{Au}$ = 0.288 nm is the diameter of the gold atom at room temperature. $r_A$, $r_B$, and $r_C$ define the major equatorial, minor equatorial, and polar radii of the triaxial ellipsoidal fit of the AuNPs[13] as noted in Figure S3, Table S1, and Table S2. The volume of an AuNP ($V_{NP}$) is calculated and follows:

$$V_{NP} = \frac{4\pi}{3} r_A r_B r_C \tag{5}$$

The electron density for AuNPs ($\rho_{NP}$) is then obtained as follows, with the number of electrons in a gold atom ($N_e$):

$$\rho_{NP} = \frac{N_e}{V_{Au}} \frac{(r_A - d_{Au})(r_B - d_{Au})(r_C - d_{Au})}{r_A r_B r_C} \quad (6)$$

Finally, with the electron density of a single gold atom $\rho_{Au} = N_e/V_{Au}$, Equation 6 simplifies to the following, which can be used to define the electron density for any ellipsoidal or spherical AuNP assuming a face-centered cubic (FCC) unit cell:

$$\rho_{NP} = \rho_{Au} \prod_{i=A,B,C} (1 - \frac{d_{Au}}{r_i}) = 3519.08\ e^-/nm^3 \quad (7)$$

<u>(GT)$_n$ ssDNA</u>

The volume for repeating units of (GT)$_n$ was estimated using the volume modeling feature in UCSF Chimera[28] and the number of electrons in the system was calculated directly by elemental composition.

#### Triaxial Ellipsoidal Fitting

Triaxial ellipsoidal fittings of AuNP scattering curves were performed using SasView by incorporating the classical physical properties of the light scattering of ellipsoidal particles in solution.[13] Here, the scattering of randomly ordered particles in solution is expressed as follows:

$$I(q) = scale(\Delta\rho)^2 \frac{V_{NP}}{4\pi} \int_{\Omega} \phi^2(qr)\, d\Omega + background \quad (8)$$

Where the integral sums all possible rotational orientations in solution, $V_{NP}$ is defined in Equation 5, and $\Delta\rho$ is the scattering length density difference between the sample and buffer. For the triaxial ellipsoidal fittings in this work, the buffer was 0.1X PBS, with a scattering length density close to $\rho_{water}$ at room temperature (9.44 x $10^{-6}$ Å$^{-2}$) used as the starting value of the fitting. Other fit parameters were left unconstrained and the DREAM algorithm[29] was used to fit the data. The fitting was further optimized by allowing for the polydispersity of the major equatorial radius ($r_A$) to be modeled as a Gaussian distribution, where the polydispersity index, PDI = standard deviation / mean (Figure S3C). Fittings are shown in Figure S3B and data are tabulated in Table S1 and Table S2.

#### ssDNA-AuNP-SWCNT Synthesis

Two alternative methods were attempted to suspend SWCNTs with ssDNA-AuNPs:

(1) **Biotin-streptavidin binding method**: Based on previous work,[30] SWCNTs (0.2 mg, NanoIntegris) were suspended with 3'-biotinylated (GT)$_6$ and (GT)$_{15}$ oligomers (1 mg,

Integrated DNA Technologies) in 1 mL total volume of 0.1X PBS via the same probe-tip sonication protocol as described earlier. Sonication was followed by centrifugation at 16,100 rcf for 30 min and separation of the supernatant containing the ssDNA-SWCNT product. The ssDNA-SWCNT solution was concentrated by centrifugal filtration (100 kDa MWCO, Millipore Sigma) with one Milli-Q water wash step. The ssDNA-SWCNT suspension was then mixed at various ratios with streptavidin-functionalized AuNPs (5 nm diameter, Cytodiagnostics Inc.). Although the mixtures were still near-infrared fluorescent (indicating biotin-ssDNA suspension of SWCNTs), no complexes of ssDNA-AuNP-SWCNTs were observed via SAXS measurement. This method was also not pursued further due to the multivalency of streptavidin (each streptavidin molecule has four attachment points for biotin), resulting in the potential for SWCNT bridging and subsequent artifacts in analysis.

(2) **Corona exchange method**: Based on our previous work,[31] SWCNTs (0.2 mg, NanoIntegris) were suspended with untagged $(GT)_{15}$ oligomers (1 mg, Integrated DNA Technologies) in 1 mL total volume of 0.1X PBS via the same probe-tip sonication protocol as described earlier. Sonication was followed by centrifugation at 16,100 rcf for 30 min and separation of the supernatant containing the ssDNA-SWCNT product. The ssDNA-SWCNT solution was concentrated, and free ssDNA was removed by centrifugal filtration (100 kDa MWCO, Millipore Sigma) with five Milli-Q water wash steps. Exchange of the initial corona (untagged ssDNA) for the final corona (AuNP-tagged ssDNA) was attempted either by passive exchange or by dialysis. For the former case, $(GT)_{15}$-AuNPs were mixed with $(GT)_{15}$-SWCNTs in molar ratios ranging from 30-30,000 AuNP-tagged ssDNA to SWCNTs (10 total ratios tested, all in 1X PBS). This range of molar ratios encompasses and far exceeds the expected value of 140 $(GT)_{15}$ strands per SWCNT,[31] in the aim of providing a driving force for AuNP-tagged ssDNA adsorption onto SWCNTs. For the latter case of dialysis-based exchange, the $(GT)_{15}$-SWCNT suspension (60 mg/L) was mixed with free $(GT)_{15}$-AuNPs (74 nM of 5 nm diameter AuNPs or 132 nM of 7 nm diameter AuNPs) and overnight dialysis (~16 h against 2 L of Milli-Q water) was used to remove untagged, desorbed ssDNA (0.5 mL volume in Slide-A-Lyzer™ Dialysis Cassettes with 20K MWCO, Thermo Fisher). For both corona exchange attempts, no complexes of ssDNA-AuNP-SWCNTs were observed via SAXS measurement. This is most likely due to the lower exchange rate for the bulkier AuNP-tagged ssDNA onto the SWCNT surface in place of the smaller untagged ssDNA already on the surface.

Purification of resulting ssDNA-AuNP-SWCNT complexes was also attempted. The centrifugation step normally included after ssDNA-SWCNT suspension (16,100 rcf for 30-90 min to remove any SWCNTs not suspended by ssDNA in the pellet) caused complete sample sedimentation due to the large AuNP mass. Centrifugal filtering was also attempted (3 kDa MWCO, Millipore Sigma). However, the combination of AuNPs and SWCNTs led to complete and irreversible sample embedding within the membrane filter.

Finally, we explored the possibility of iron nanoparticle contamination in SWCNT samples accounting for the X-ray scattering results. Namely, the HiPco SWCNT synthesis process involves the *in situ* thermal decomposition of iron pentacarbonyl to iron catalyst nanoclusters upon which the SWCNTs then grow.[32] Raw HiPco SWCNTs are reported to contain <35 wt% residual iron catalyst (NanoIntegris characterization data). We have previously measured approximately 8.5 wt% iron present in our raw HiPco SWCNT samples once suspended with ssDNA, via X-ray photoelectron spectroscopy analysis (unpublished). To test the potential influence of iron nanoclusters, we further tested super-purified HiPco SWCNTs (NanoIntegris; reported to contain <5 wt% residual iron catalyst and presumably also less once ssDNA-suspended) similarly suspended in ssDNA. Baseline X-ray scattering at the relevant SWCNT concentration with these super-purified samples was negligible in comparison to that in the presence of AuNPs, supporting the insignificant contribution of residual iron catalyst in the observed scattering profiles.

#### *Ab initio* modeling of ssDNA-AuNP-SWCNT complexes

We adapt an *ab initio* modeling technique to produce 3D models complementing our 2D description of the AuNP geometries on the SWCNT surface. A large parameter space of initialization geometries was explored to converge on best-fit *ab initio* models for $(GT)_{15}$- and $(GT)_{6}$-AuNP-SWCNT complexes. In structural biology, SAXS profiles from macromolecules are traditionally modeled via *ab initio* shape reconstruction using software such as DAMMIN[33], GASBOR[34], or more recently, DENSS[35]. However, these 3D modeling techniques all fail to capture samples with very high aspect ratios, such as SWCNTs. 3D modeling of the AuNPs adsorbed on the SWCNT surface was instead accomplished using the *ab initio* modeling capabilities of SASHEL, expanding upon the methodology described by its developers Burian and Amenitsch.[25] SASHEL was adapted to move clusters of 1000 dummy atoms, where each cluster represents a single AuNP, starting from curated initial geometries (see SI Methods and Figure S16A). The best-fit models obtained using this technique demonstrate the dynamic nature of the system, providing a larger graphical view of the system as it exists in solution and complementing our statistical analysis of inter-AuNP distances.

It was initially determined that both $(GT)_{15}$- and $(GT)_{6}$-AuNP-SWCNTs were best captured by parallel lines of AuNPs (SWCNTs) starting at a distance of 15 nm apart (Table S4 and Figure S17). Based on our statistical analysis, the average radial inter-AuNP distance for $(GT)_{6}$-AuNP-SWCNTs is 7.6 ± 0.6 nm. Thus, *ab initio* modeling results demonstrate distances over two-fold higher, indicating that two neighboring ssDNA-AuNP-SWCNTs are required to properly describe the overall volume of the electron density of the samples.

The starting stack-building block heights ($H_{BB}$) and hence the initial number of AuNPs ($N_{NP}$) ranging from 2 to 18 AuNPs were attempted in order to expand the volume of the modeling space and modulate complexity within the modeling (Table S5, Figure S18, and Figure S19). The $H_{BB}$ range attempted represent integer multiples of the inter-AuNP block heights ($H_{GB}$) for $(GT)_{15}$- and $(GT)_6$-AuNP-SWCNT (14.3 and 17.2 nm, respectively), as determined by the average axial inter-AuNP distances from *P(r)* functions. Both $(GT)_{15}$ and $(GT)_6$-AuNP-SWCNTs demonstrated local minima in $\chi^2$-values when plotted as a function of $H_{BB}$, with average minima at integer multiples of 54.5 ± 2.8 nm (Figure S20). This $H_{BB}$ value could represent the upper end of the observed maximum length dimension, $d_{max}$, range of the ssDNA-AuNP-SWCNT samples (~45-55 nm) as determined from *P(r)* functions (Figure 1A-B). $(GT)_{15}$ and $(GT)_6$-AuNP-SWCNTs were best fit using starting models Containing 16 AuNPs with $H_{BB}$ of 114.5 nm and 18 AuNPs with $H_{BB}$ of 155.2 nm, respectively (Table S5 and Figure S21A-B). Initially, these AuNPs are evenly arranged according to experimental geometries (see SI Methods and Figure S16), leading to calculated AuNP packing of 0.14 and 0.12 AuNP per nm for $(GT)_{15}$ and $(GT)_6$-AuNP-SWCNT, respectively. While the calculated AuNP packing of $(GT)_{15}$-AuNP-SWCNT complemented those calculated from TEM analysis (0.139 AuNPs per nm length of SWCNT), those calculated for $(GT)_6$-AuNP-SWCNT were inconsistent with the corresponding 0.185 AuNPs per nm length of SWCNT from TEM analysis. Interestingly, the results of the *ab initio* modeling for $(GT)_{15}$-AuNP-SWCNTs maintain a consistent number of AuNPs, while for $(GT)_6$-AuNP-SWCNTs some dummy atoms split off from the original AuNPs and start to form new AuNPs, suggesting that there are regions of missing electron density that necessitate additional AuNPs. For purposes of comparison, the single ssDNA-AuNP-SWCNT showing the most clearly defined AuNPs was isolated from each model and compared to theoretical models produced using Solidworks (Dassault Systèmes) directly from geometries obtained from statistical analysis of the *P(r)* functions (Figure S21C). As predicted, the isolated $(GT)_{15}$-AuNP-SWCNT model shows a resemblance to the corresponding theoretical model in number and placement of AuNPs along the SWCNT axis, while the $(GT)_6$-AuNP-SWCNT model shows clear regions of missing electron density. The initial $N_{NP}$ was doubled for the starting $(GT)_6$-AuNP-SWCNT model (Figure S16C) leading to a slight reduction in $\chi^2$-value and more consistent AuNPs (dummy-atom clusters) as shown in Figure S13D and Figure S21A. The best single SWCNT was again isolated from this revised double SWCNT model and compared against the corresponding theoretical model. The results were a clear improvement in number and placement of AuNPs along the SWCNT axis (Figure S21D). To confirm the *ab initio* modeling results, the two best starting models were switched for $(GT)_{15}$- and $(GT)_6$-AuNP-SWCNTs, resulting in a lower quality of fit in both cases (Figure S22) and supporting the ssDNA sequence-specific modeling outcomes.

It was realized that 1000 dummy atoms per AuNP was insufficient to properly model the ssDNA-AuNP-SWCNT systems resulting in the software compensating for lack of electron density by expanding the model widths instead of lengths. Thus, to further explore whether the system could be modeled as a single SWCNT, the best fit double SWCNT model for $(GT)_{15}$- and $(GT)_6$-AuNP-SWCNT was selected (Figure S21) and the single SWCNT region showing the clearest AuNP spacing was extracted (Figure S21C-D). Over many iterations, the positions and density (number of dummy atoms) of the AuNPs were carefully modified leading to a best fitting single SWCNT models (see SI Methods and Figure S13A-C). The $\chi^2$-value for the best single SWCNT model is far higher than the best double SWCNT model ($\chi^2$ = 1.67 vs 0.20). Upon closer inspection of the data, the residuals of the fit show that the single SWCNT models fit better at lowest q-values (Guinier region, defining the overall size) but worse in the midrange q-values (Porod region, defining the volume and morphology) than that of the double SWCNT model (Figure S13C). This observation suggests that the increased volume requirement may eventually be overcome with a sufficiently elongated and complex (more dummy atoms) model.

### Supplementary Figures and Tables

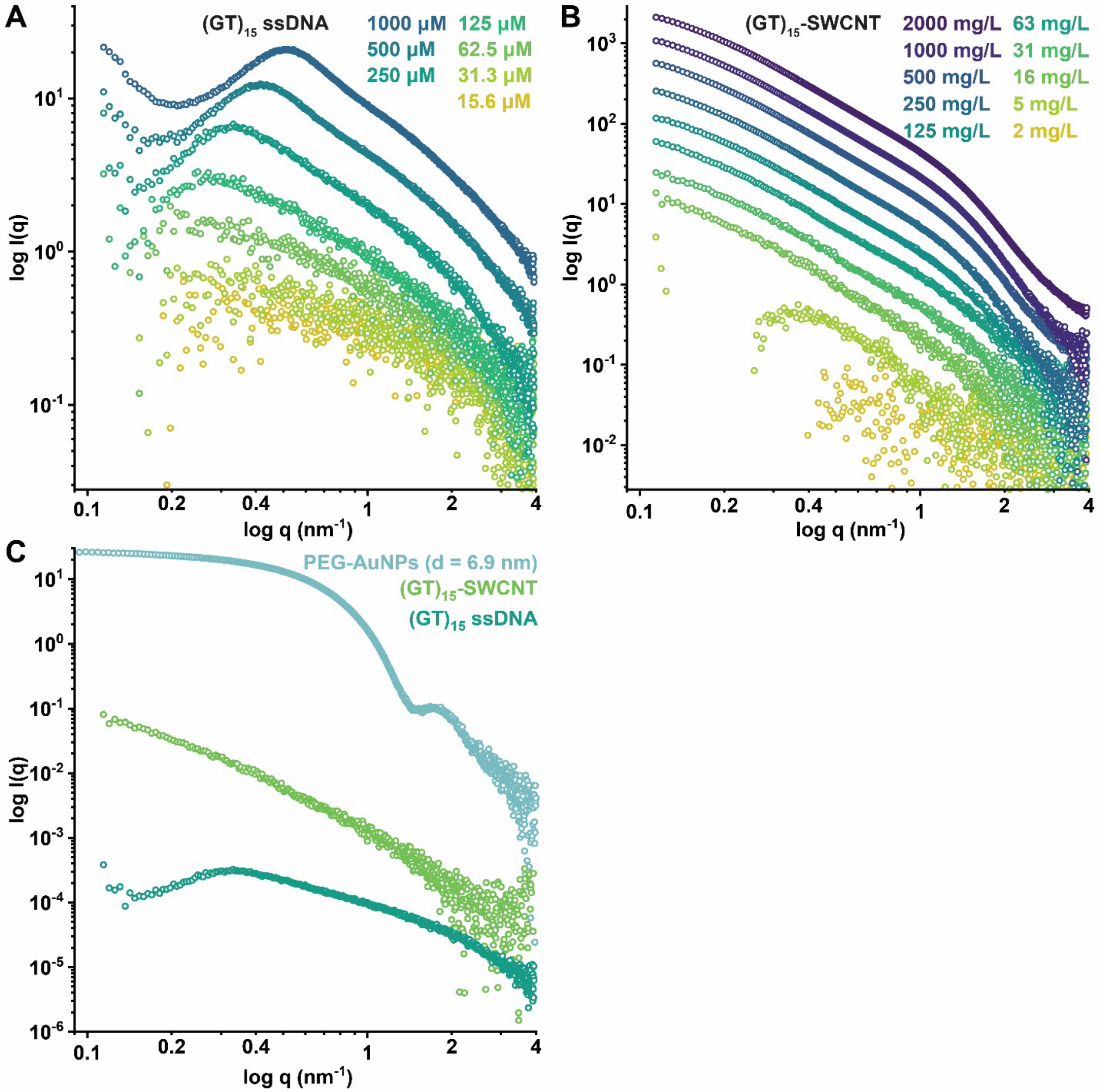


**Figure S1.** Scattering profiles as a function of concentration for serial dilutions of (A) $(GT)_{15}$ ssDNA alone and (B) $(GT)_{15}$-SWCNTs. (C) Absolute-scale scattering measurements for representative samples of 6.9 nm diameter PEG-AuNP, $(GT)_{15}$-SWCNTs, and $(GT)_{15}$ ssDNA alone, at the correct relative concentrations (i.e., 250 nM AuNP and ssDNA per 1 mg/L SWCNT).

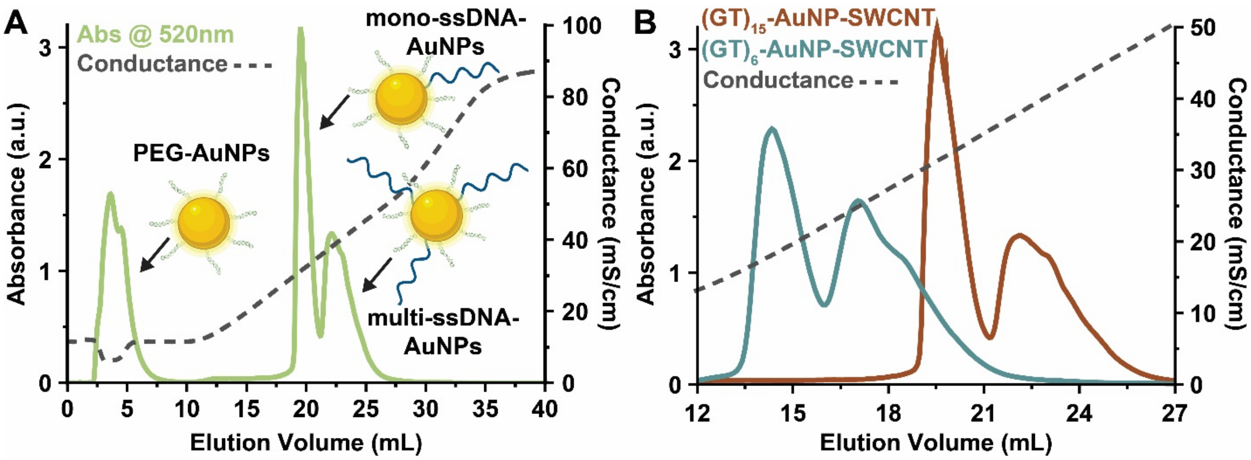


**Figure S2.** Anion exchange chromatograms for (A) the separation of non-conjugated PEG-AuNPs, mono-conjugated $(GT)_{15}$-AuNP, and multi-conjugated $(GT)_{15}$-AuNP using fast protein liquid chromatography (FPLC) and (B) $(GT)_{15}$- (red) vs. $(GT)_6$-AuNP (blue), demonstrating the length-dependent shift in the elution volume. Spectra are measured by diode array detector (DAD) at 520 nm. Conductance measurements depict salt gradient conditions (dash grey).

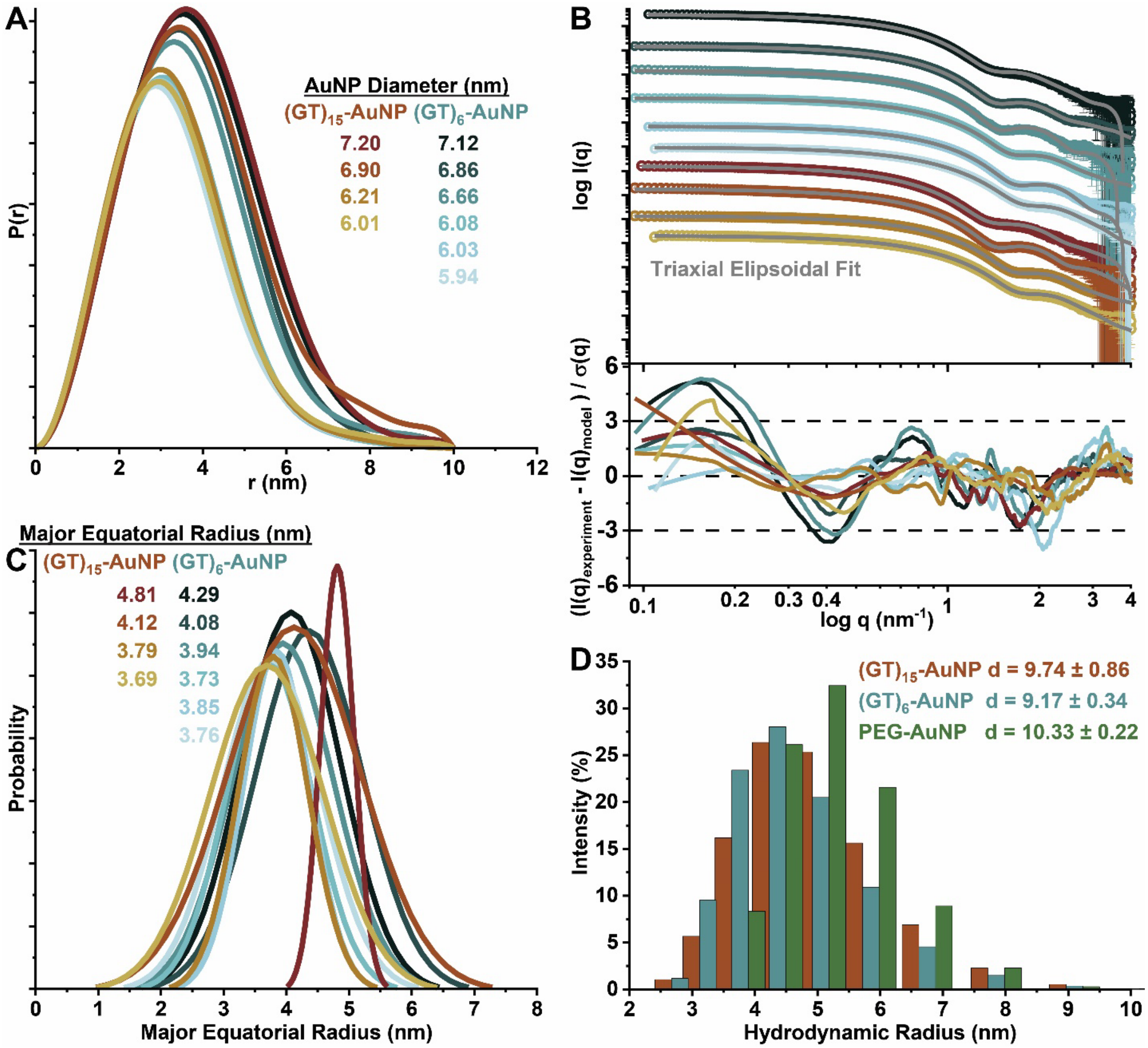


**Figure S3.** SAXS analysis for synthesized ssDNA-AuNPs. (A) Pairwise distribution functions, *P(r)*, for AuNPs of various sizes, scaled to the calculated average diameters for visual clarity. (B) Experimental SAXS profiles with calculated triaxial ellipsoidal fits (grey) for the prepared ssDNA-AuNPs (top) and the fit residuals (bottom). Scattering curves are offset for clarity and colored as in panel (A). Numerical values are summarized in Table S1 and Table S2. (C) Polydispersity of the major equatorial radius ($r_A$) modeled as a Gaussian distribution using SasView. Plots are scaled to the calculated $r_A$ values and colored as in panel (A). Numerical values for panels (A-B) are summarized in Table S1 and Table S2. (D) DLS histograms reveal increased hydrodynamic radii of AuNPs after $(GT)_n$ ssDNA conjugation and mPEG-SH coating, as compared to diameters calculated from corresponding *P(r)* functions (Table S3). Note that all ssDNA-AuNPs are also PEGylated.

**Table S1.** Physical parameters of synthesized $(GT)_6$-AuNPs obtained from pairwise distribution functions and triaxial ellipsoidal fits of scattering curves.

| ***P(r)* Peak Diameter (nm)** | **Major Equatorial Radius, $r_A$ (nm)** | **Minor Equatorial Radius, $r_B$ (nm)** | **Polar Radius, $r_C$ (nm)** | **Polydispersity Index (PDI)** | **$\chi^2$-value** | **$r_A$ / $r_C$** |
|---|---|---|---|---|---|---|
| 5.94 ± 0.09 | 3.76 ± 0.08 | 2.88 ± 0.04 | 2.07 ± 0.01 | 0.21 | 0.74 | 1.8 |
| 6.03 ± 0.07 | 3.85 ± 0.03 | 2.86 ± 0.02 | 2.25 ± 0.01 | 0.14 | 2.48 | 1.7 |
| 6.08 ± 0.03 | 3.73 ± 0.16 | 3.08 ± 0.05 | 2.24 ± 0.04 | 0.18 | 0.88 | 1.7 |
| 6.66 ± 0.04 | 3.94 ± 0.12 | 3.26 ± 0.18 | 2.61 ± 0.16 | 0.2 | 2.41 | 1.5 |
| 6.86 ± 0.04 | 4.08 ± 0.03 | 3.43 ± 0.01 | 2.60 ± 0.01 | 0.19 | 0.66 | 1.6 |
| 7.12 ± 0.06 | 4.29 ± 0.04 | 3.65 ± 0.02 | 2.60 ± 0.01 | 0.2 | 1.96 | 1.7 |

**Table S2.** Physical parameters of synthesized $(GT)_{15}$-AuNPs obtained from pairwise distribution functions and triaxial ellipsoidal fits of scattering curves.

| ***P(r)* Peak Diameter (nm)** | **Major Equatorial Radius, $r_A$ (nm)** | **Minor Equatorial Radius, $r_B$ (nm)** | **Polar Radius, $r_C$ (nm)** | **Polydispersity Index (PDI)** | **$\chi^2$-value** | **$r_A$ / $r_C$** |
|---|---|---|---|---|---|---|
| 6.01 ± 0.08 | 3.69 ± 0.08 | 2.97 ± 0.03 | 2.15 ± 0.01 | 0.25 | 1.21 | 1.7 |
| 6.21 ± 0.08 | 3.79 ± 0.12 | 2.97 ± 0.06 | 2.20 ± 0.01 | 0.15 | 0.7 | 1.7 |
| 6.90 ± 0.02 | 4.12 ± 0.11 | 3.39 ± 0.03 | 2.65 ± 0.03 | 0.26 | 0.33 | 1.6 |
| 7.20 ± 0.02 | 4.81 ± 0.15 | 3.48 ± 0.09 | 2.64 ± 0.02 | 0.06 | 1.1 | 1.8 |

**Table S3.** Physical parameters of synthesized AuNPs as obtained from pairwise distribution functions of X-ray scattering and dynamic light scattering.

| | ***P(r)* Peak Diameter (nm)** | **DLS Peak Diameter (nm)** | **Increase in Diameter from PEGylation (nm)** |
|---|---|---|---|
| $(GT)_{15}$-AuNP | 6.90 ± 0.02 | 9.74 ± 0.86 | 2.83 ± 0.88 |
| $(GT)_6$-AuNP | 6.86 ± 0.04 | 9.17 ± 0.34 | 2.31 ± 0.38 |
| PEG-AuNP | 6.82 ± 0.03 | 10.33 ± 0.22 | 3.51 ± 0.25 |

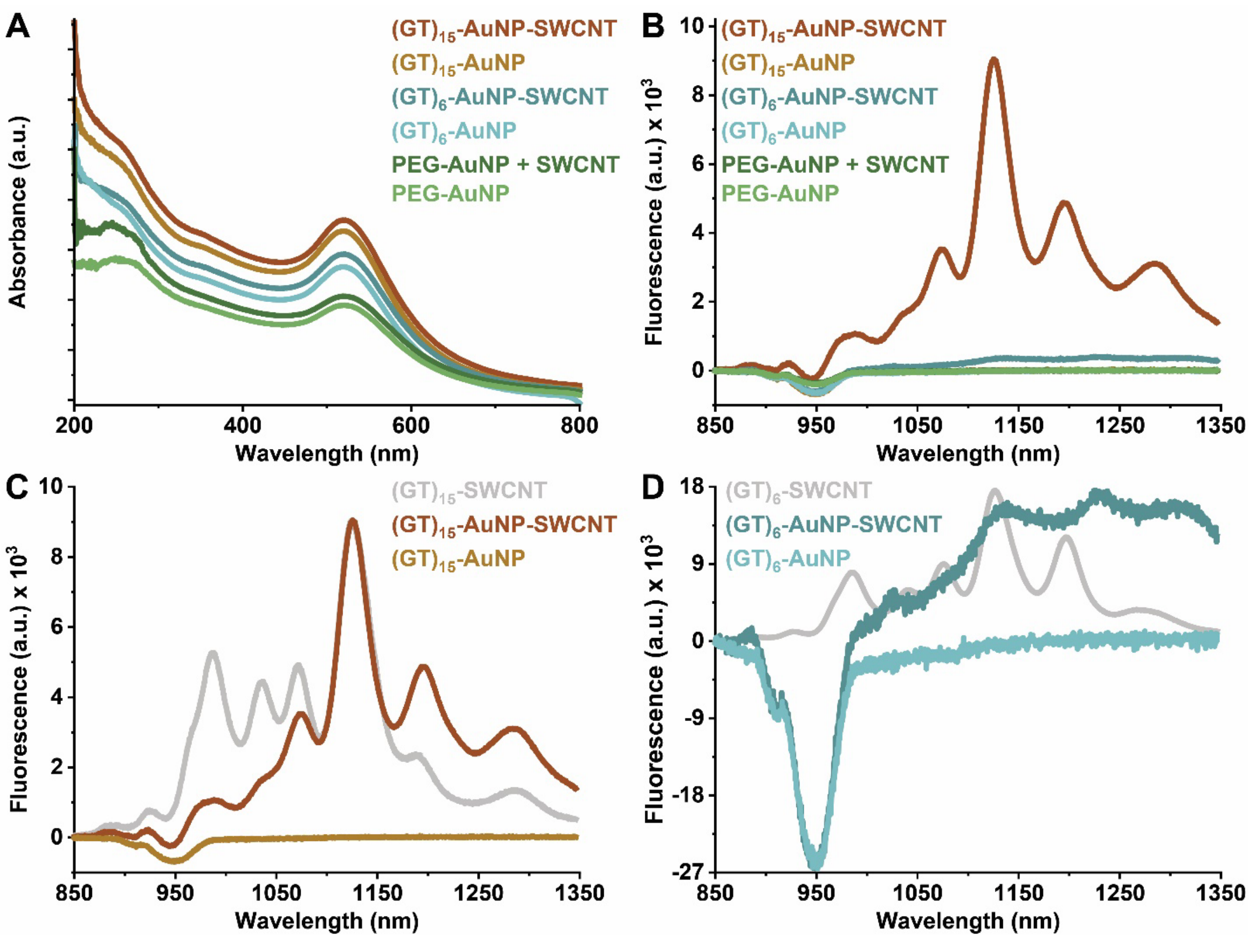


**Figure S4.** ssDNA-AuNP-SWCNT optical characterization. (A) Absorbance spectra for $(GT)_{15}$-AuNP alone and suspending SWCNTs (orange and red, respectively), $(GT)_6$-AuNP alone and suspending SWCNTs (light and dark blue, respectively), and PEG-AuNP alone and attempted-to-suspend SWCNTs (light and dark green, respectively). The consistent AuNP plasmon resonance peak at approximately 520 nm demonstrates that AuNP tags are intact after the probe-tip sonication suspension process with SWCNTs. Spectra are offset for clarity. Note that these SWCNT concentrations of approximately 0.2 mg/L produce negligible near-infrared absorbance fingerprints and this region of the spectrum is therefore omitted. (B) Fluorescence spectra for the same sample set as panel (A) confirm SWCNT suspension with the presence of near-infrared fluorescence. (C) $(GT)_{15}$-AuNP-SWCNT and $(GT)_{15}$-SWCNT samples are compared to a $(GT)_{15}$-SWCNT conjugate without AuNP tags, with peaks normalized to maximum emission intensity. (D) $(GT)_6$-AuNP-SWCNT and $(GT)_6$-SWCNT samples are compared to a $(GT)_6$-SWCNT conjugate without AuNP tags, with peaks normalized to maximum emission intensity. All AuNPs are synthesized diameters of 5.9-7.2 nm. All fluorescence measurements were obtained with 721 nm laser excitation.

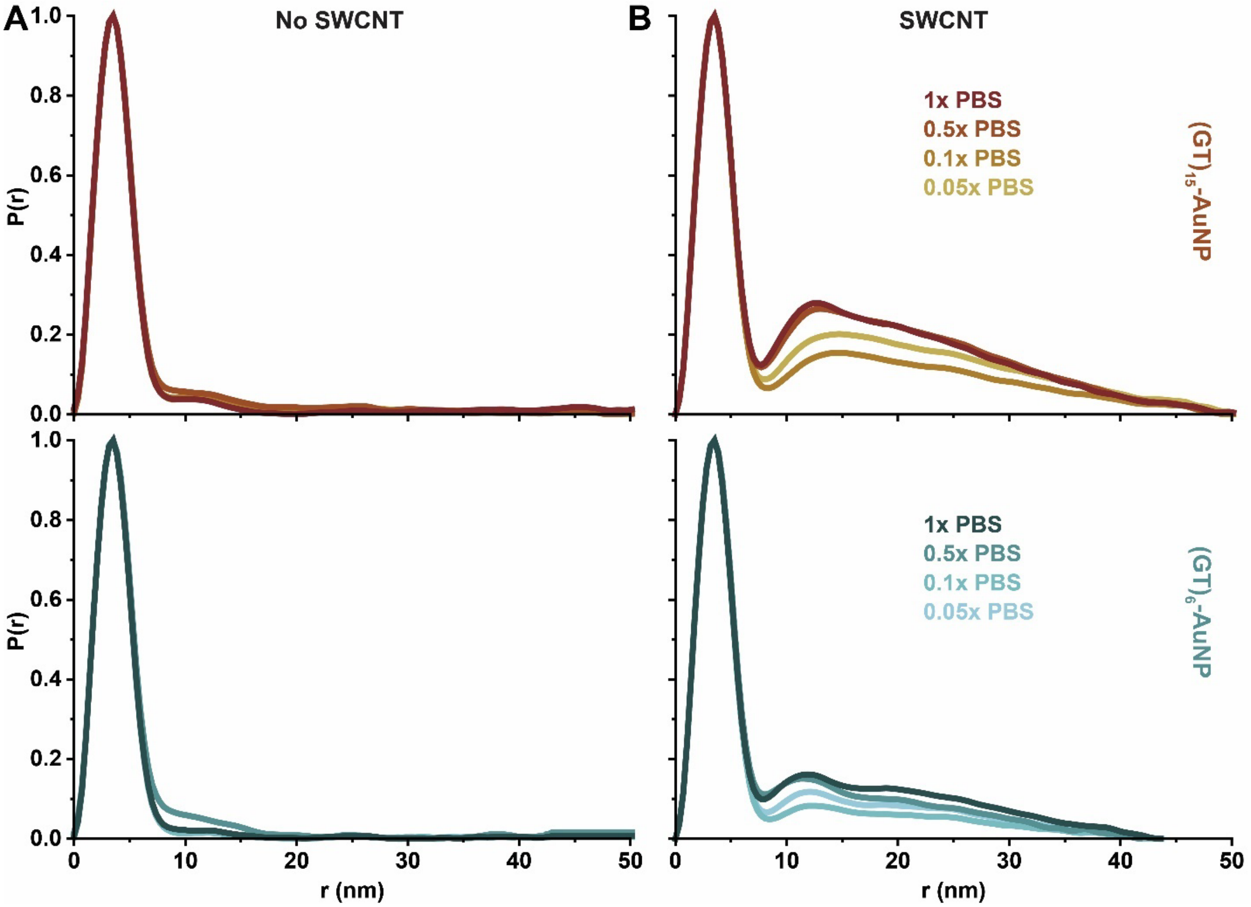


**Figure S5.** Representative pairwise distribution functions, *P(r)*, of AuNP-tagged ssDNA either (A) free in solution or (B) adsorbed to SWCNTs, as a function of ionic strength over a range of 0.05X to 2X PBS or corresponding Debye lengths, $\lambda_D$ = 3.37-0.53 nm. (Top) $(GT)_{15}$-AuNPs vs. $(GT)_{15}$-AuNP-SWCNT complexes and (bottom) $(GT)_6$-AuNPs vs. $(GT)_6$-AuNP-SWCNT complexes. *P(r)* functions are normalized to the intra-AuNP peak to compensate for slight fluctuations in X-ray beam intensity or sample concentration. Order only emerges in the presence of ssDNA-AuNPs adsorbed to SWCNTs.

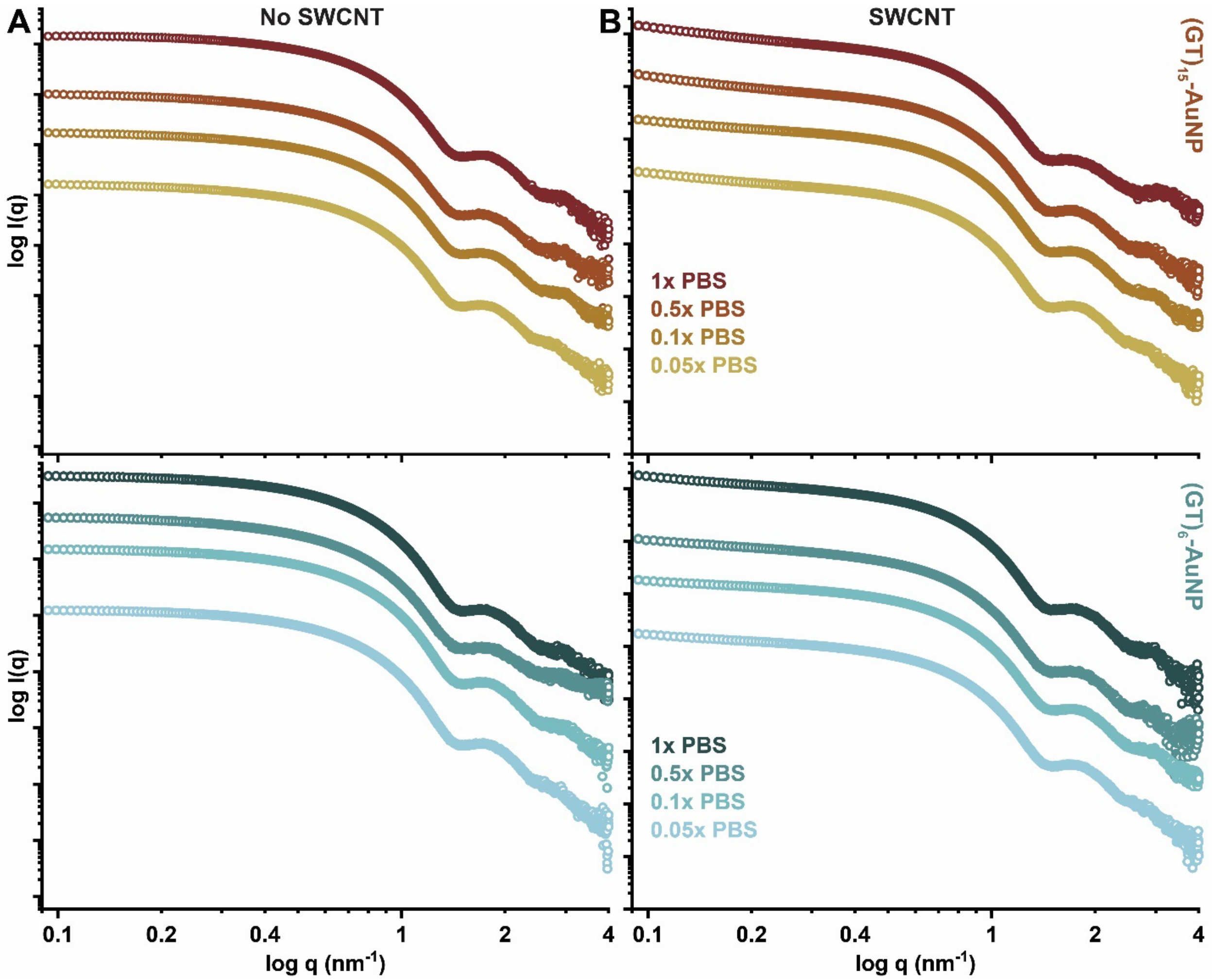


**Figure S6.** Representative scattering curves of AuNP-tagged ssDNA either (A) free in solution or (B) adsorbed to SWCNTs, as a function of ionic strength over a range of 0.05X to 2X PBS or corresponding Debye lengths, $\lambda_D$ = 3.37-0.53 nm. (Top) $(GT)_{15}$-AuNPs vs. $(GT)_{15}$-AuNP-SWCNT complexes and (bottom) $(GT)_6$-AuNPs vs. $(GT)_6$-AuNP-SWCNT complexes. Scattering curves are offset for clarity. The major difference of note in curves (A) without vs. (B) with SWCNTs is found in the lowest q-value range ($q < 0.2$ $nm^{-1}$): the upward curvature at low q with SWCNTs represents interparticle interaction from the ordering of AuNPs on the SWCNT surface.

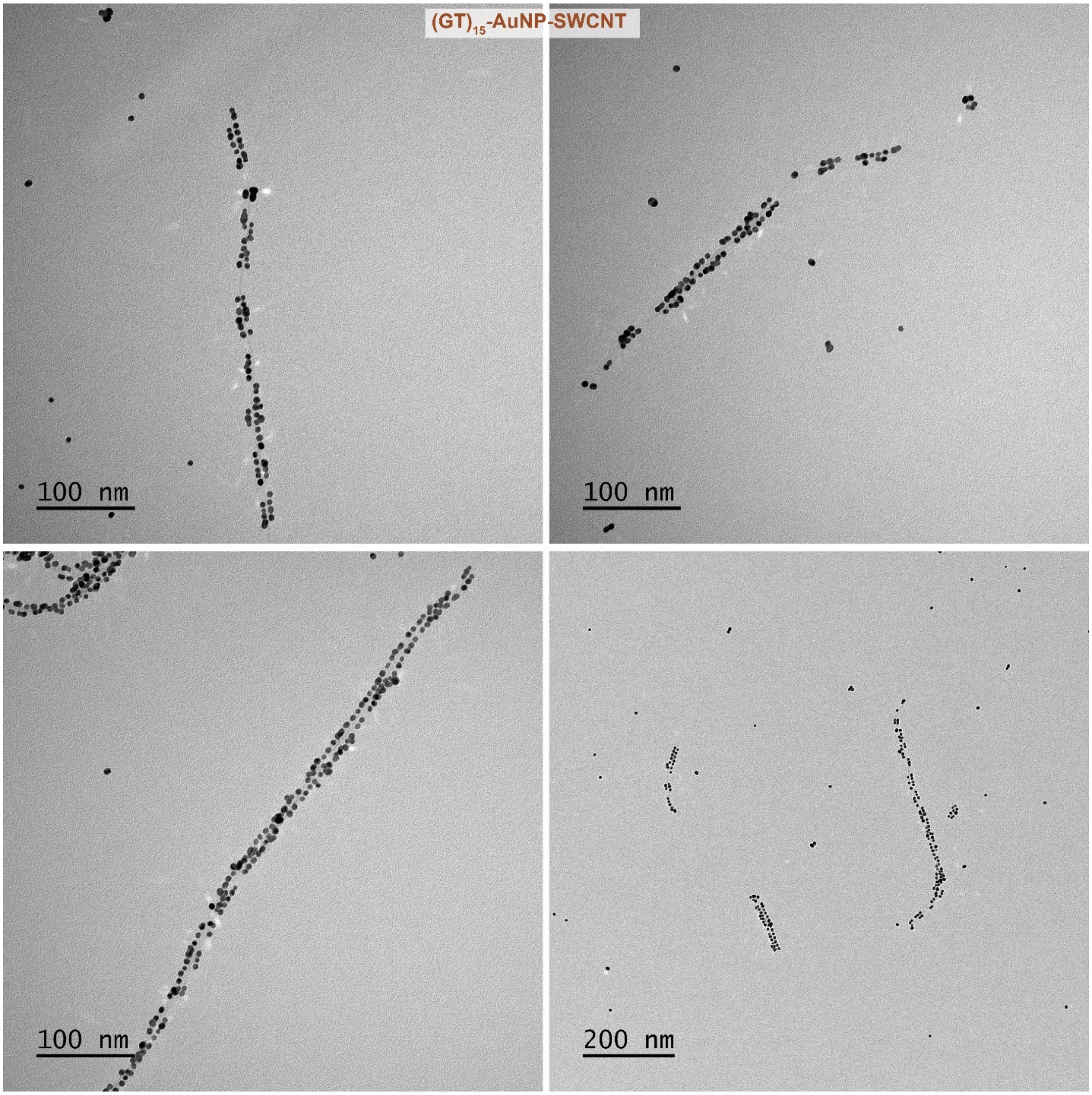


**Figure S7.** Representative TEM images for $(GT)_{15}$-AuNP-SWCNTs.

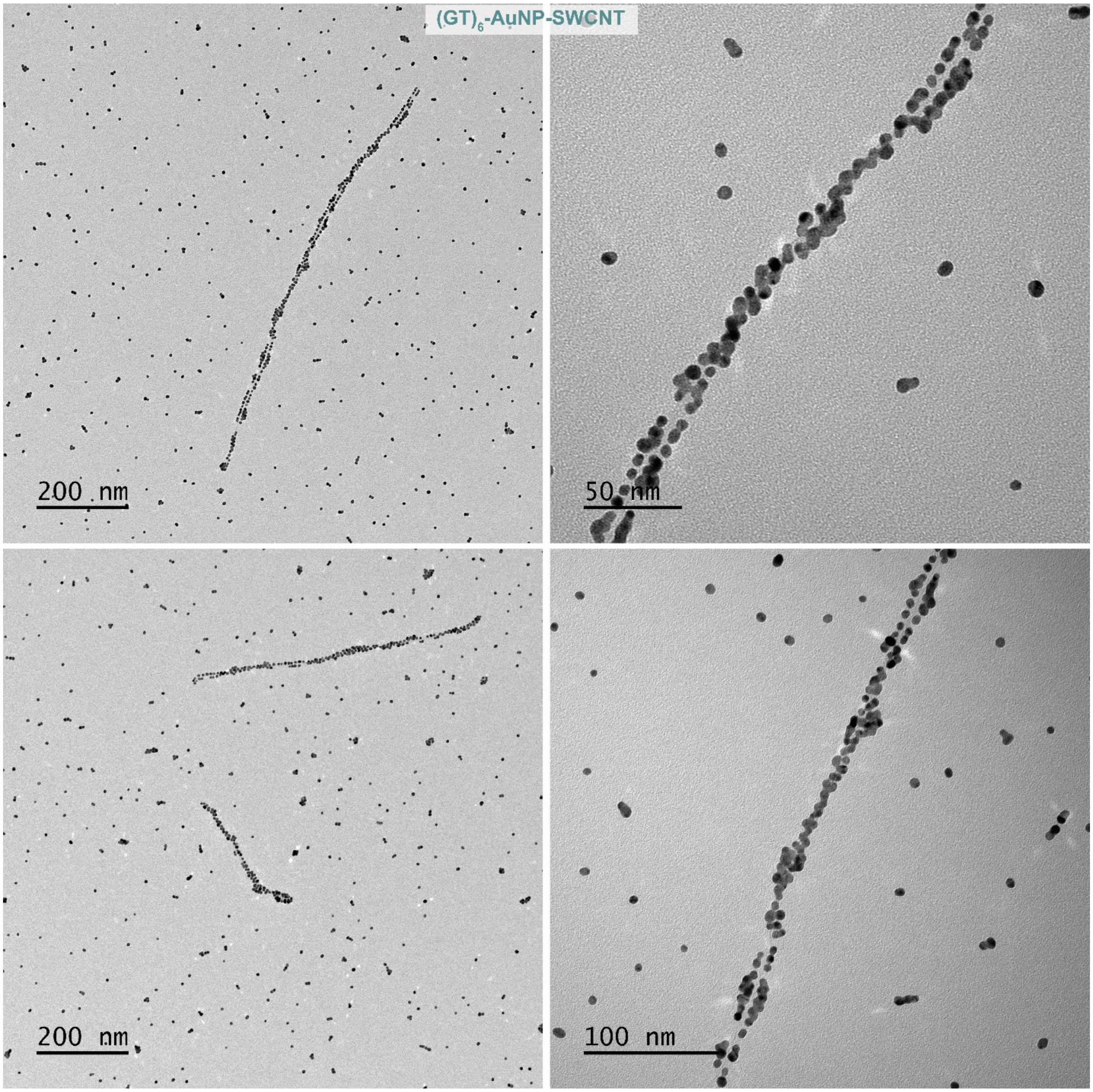


**Figure S8.** Representative TEM images for $(GT)_6$-AuNP-SWCNTs.

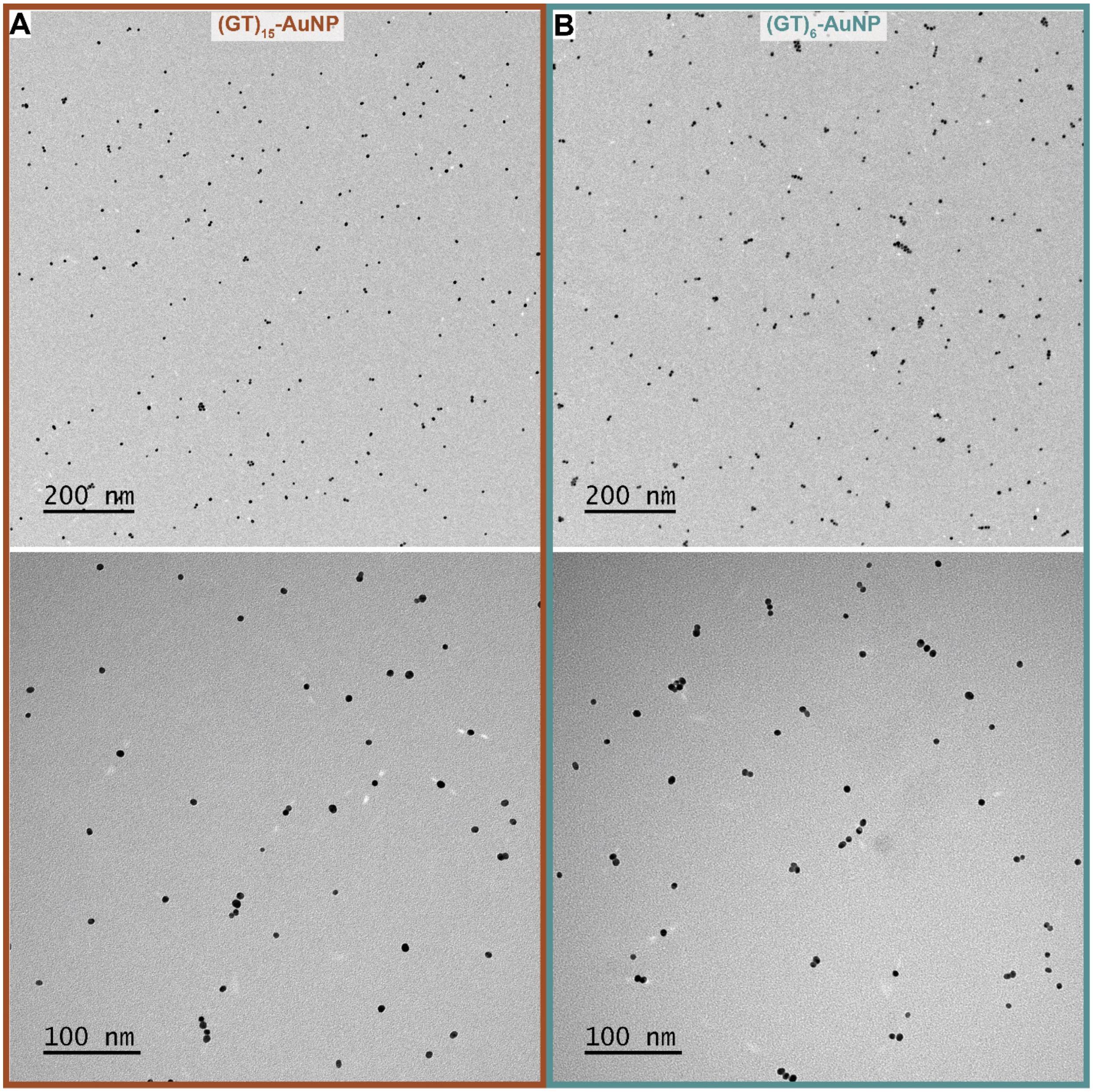


**Figure S9.** Representative TEM images for (A) $(GT)_{15}$-AuNPs and (B) $(GT)_6$-AuNPs prepared by the same method as ssDNA-AuNP-SWCNTS but absent the SWCNT substrate. All controls do not show order when free in the solution state, in the absence of SWCNTs.

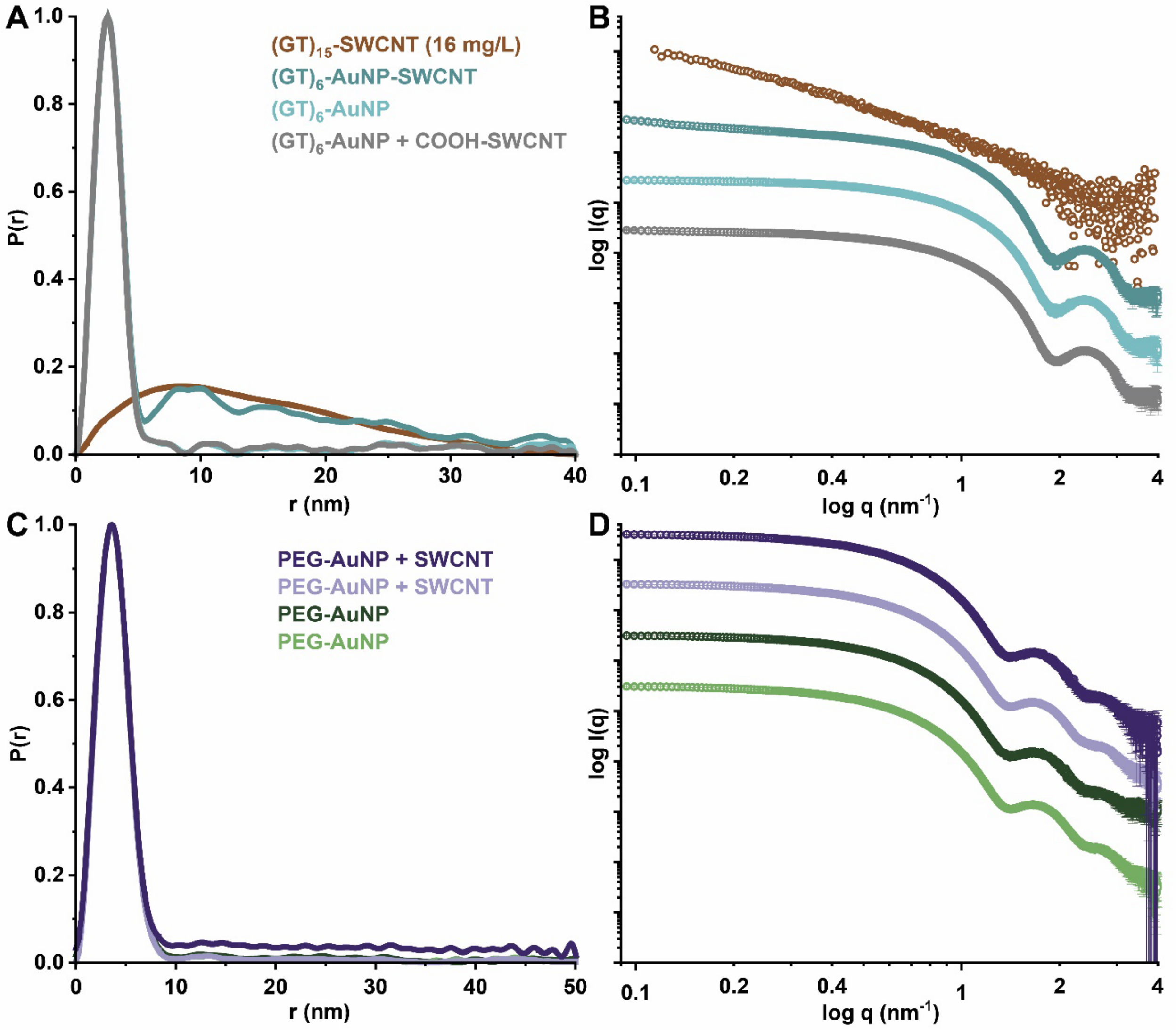


**Figure S10.** Experimental controls for ssDNA-AuNP-SWCNT preparations. (A) Pairwise distribution functions, *P(r)*, and (B) scattering curves for $(GT)_{15}$-SWCNTs (no AuNPs) at the lowest measurable concentration (16 mg/L), $(GT)_6$-AuNP-SWCNTs (full complex), $(GT)_6$-AuNPs (no SWCNTs), $(GT)_6$-AuNP-SWCNTs (2 mg/L), and $(GT)_6$-AuNPs mixed with carboxylated SWCNTs (2 mg/L; no suspension). The *P(r)* function for $(GT)_{15}$-SWCNT (no AuNPs) is normalized to the inter-AuNP peak of $(GT)_6$-AuNPs-SWCNT for clarity. (C) *P(r)* function and (D) scattering curves for two batches of PEG-AuNPs (green) vs. PEG-AuNPs attempted-to-suspend with SWCNTs (purple) by the same method as ssDNA-AuNP-SWCNTS. All AuNP samples are normalized to intra-AuNP peak.

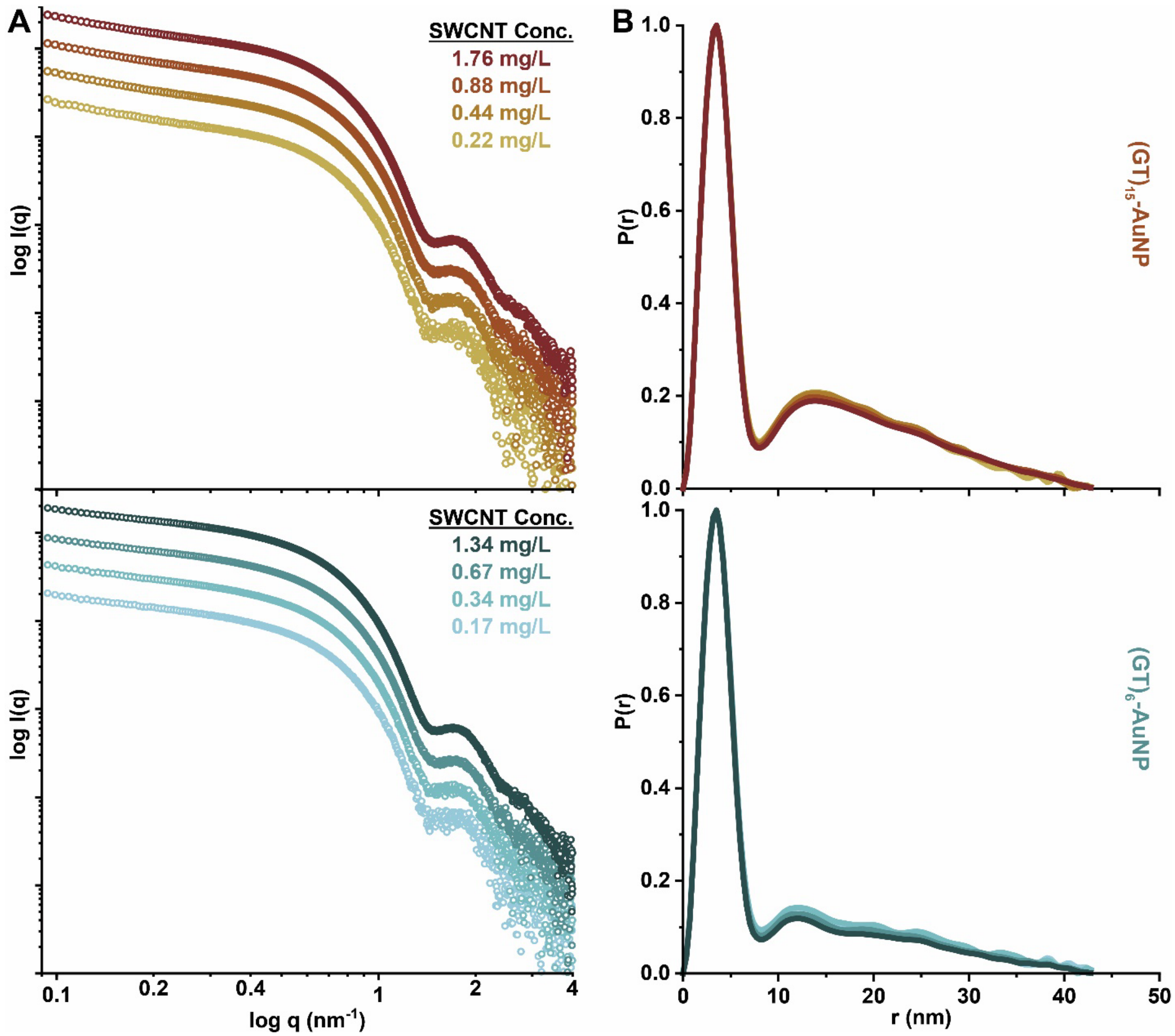


**Figure S11.** (A) Scattering curves of AuNP-tagged ssDNA adsorbed to SWCNTs, as a function of SWCNT concentration for (top) $(GT)_{15}$-AuNP-SWCNT complexes and (bottom) $(GT)_6$-AuNP-SWCNT complexes. Scattering curves are offset for clarity. (B) Pairwise distribution functions, *P(r)*, of AuNP-tagged ssDNA adsorbed to SWCNTs, as a function of concentration. *P(r)* functions are normalized to the intra-AuNP peak to compensate for fluctuations in intensity. Samples are observed over SWCNT concentrations relevant to biological applications (0.17-1.76 mg/L).

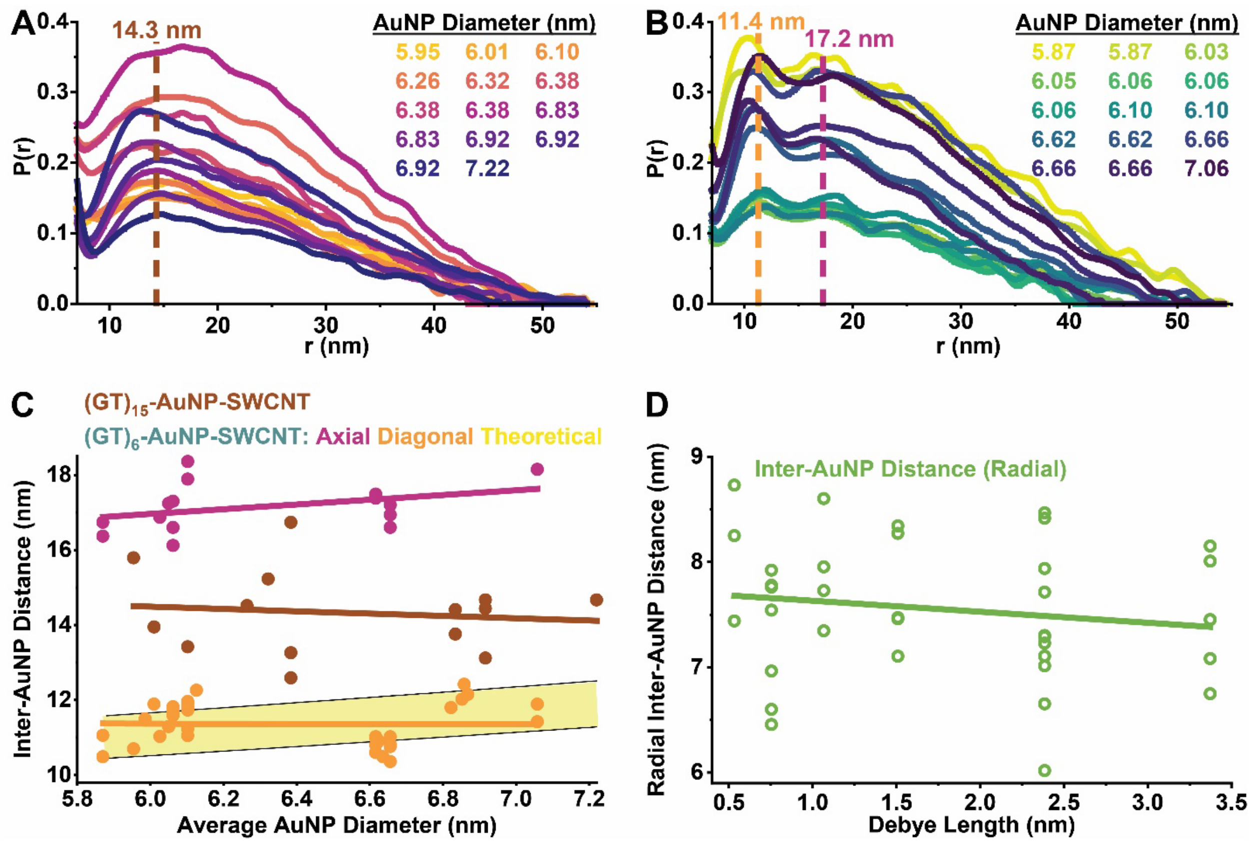


**Figure S12.** Pairwise distribution functions, *P(r)*, for samples with varying AuNP sizes ranging from 5.9 to 7.2 nm diameter for (A) $(GT)_{15}$-AuNP-SWCNTs and (B) $(GT)_6$-AuNP-SWCNTs that demonstrate a clear second inter-AuNP distance. Dashed vertical lines are added to visualize the average inter-AuNP distances for $(GT)_{15}$-AuNP-SWCNTs (red), and $(GT)_6$-AuNP-SWCNTs diagonal (orange) and axial (magenta). *P(r)* functions are normalized to the primary intra-AuNP peak, then the x-axis minimum is set to focus on the inter-AuNP peak for clarity. (C) Plots inter-AuNP distances for $(GT)_{15}$-AuNP-SWCNTs (red), and $(GT)_6$-AuNP-SWCNTs diagonal (orange) and axial (magenta) as a function of AuNP size. Only the diagonal inter-AuNP distances should be subject to changes as a function of AuNP size as displayed graphically by theoretical distances (yellow) calculated with AuNPs fixed on the SWCNT surface. (D) Plots of radial inter-AuNP distances (green) for $(GT)_6$-AuNP-SWCNTs as a function of Debye length ($\lambda_D$) as calculated from experimental diagonal and axial inter-AuNP distances from a single size of AuNPs (d = 6.1 ± 0.03 nm).

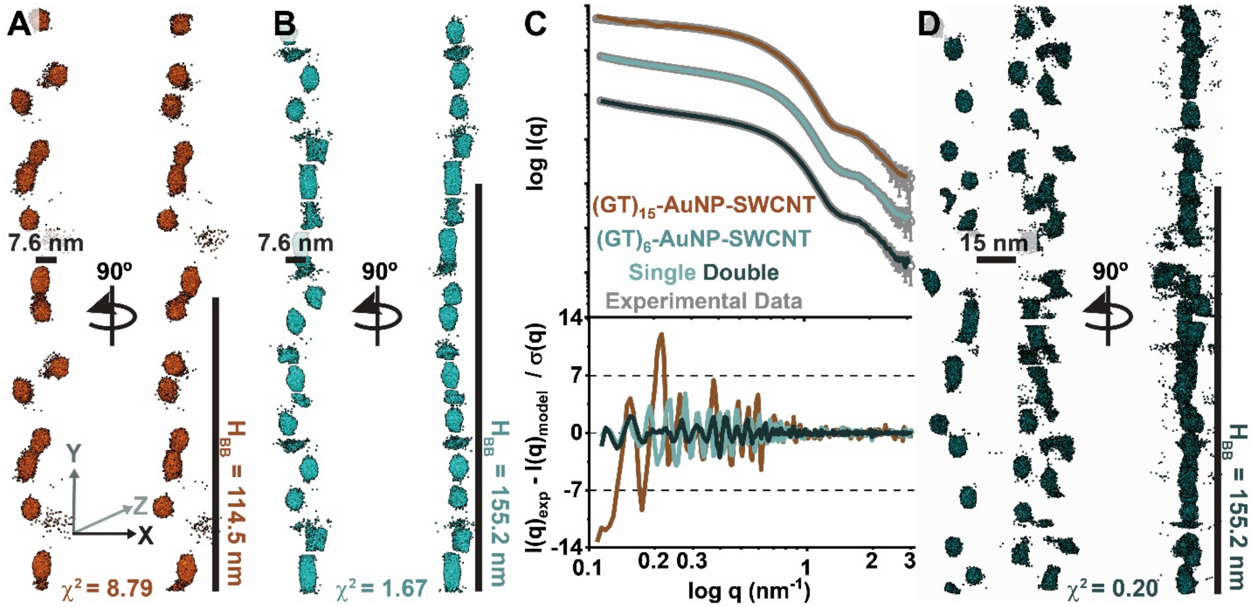


**Figure S13.** Best fit single SWCNT *ab initio* modeling results for (A) $(GT)_{15}$-AuNP-SWCNT and (B) $(GT)_6$-AuNP-SWCNTs. (C) SAXS profiles with fits and residuals for each model colored as in panel (A, B, and D). Scattering curves are offset for clarity. (D) *Ab initio* modeling results for $(GT)_6$-AuNP-SWCNTs modeled as two parallel SWCNTs for comparison. Initial models started with a stack building block height ($H_{BB}$) of 114.5 and 155.2 nm for $(GT)_{15}$- and $(GT)_6$-AuNP-SWCNT, respectively, as defined by the best fit starting model parameters found in Table S5. Final $\chi^2$-values shown beneath each model.

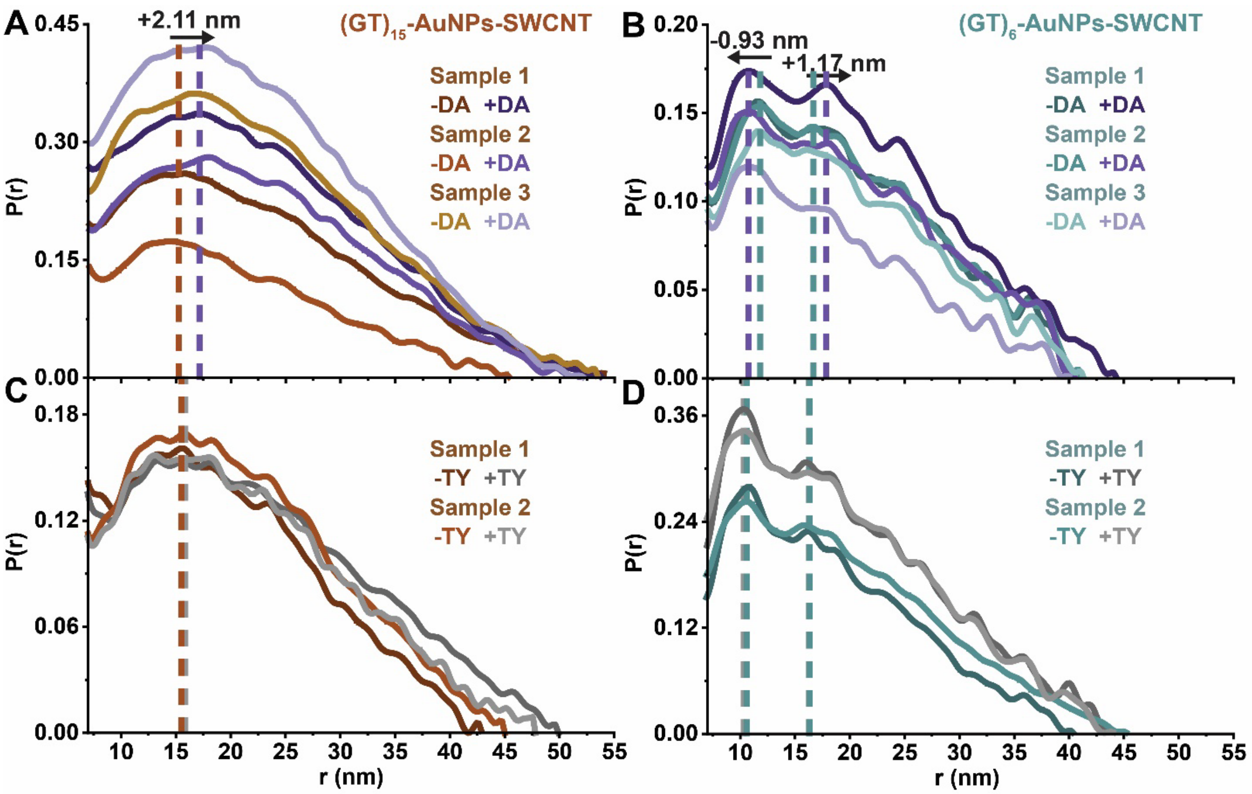


**Figure S14.** Pairwise distribution functions, *P(r)*, for all replicates alone for (A-C) $(GT)_{15}$-AuNP-SWCNTs (red-orange series) and (B-D) $(GT)_6$-AuNP-SWCNTs (blue series) or in the presence of (A-B) DA (purple series) or (C-D) TY (grey series). Dashed vertical lines are added to visualize peak shifts. *P(r)* functions are normalized to the primary intra-AuNP peak, then the x-axis minimum is set to focus on the inter-AuNP peak for clarity.

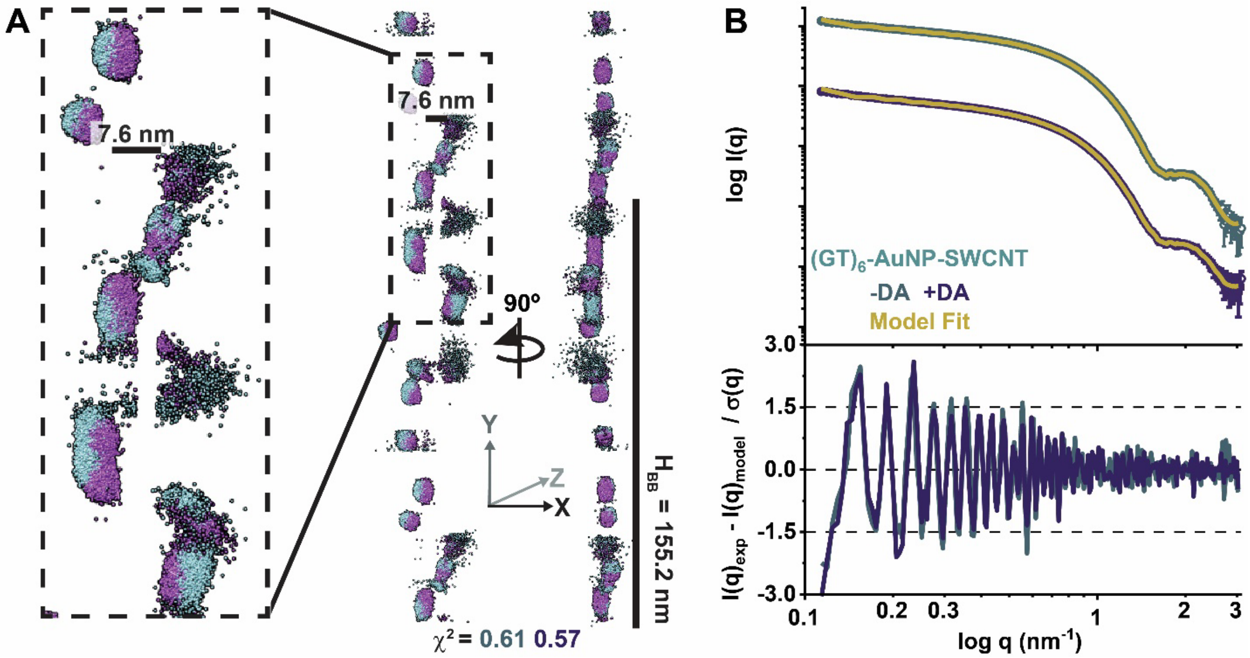


**Figure S15.** Comparison of *Ab initio* modeling results for $(GT)_6$-AuNP-SWCNTs modeled from scattering profiles with (blue series) and without dopamine (DA; purple series) Initial model started from the best single SWCNT model (Figure 1F and Figure S13C) started with two parallel rows of AuNPs set 7.5 nm apart. Both models start with a stack building block height ($H_{BB}$) of 155.2 nm and number of stacks ($N_S$) of 2 as defined by the best fit starting model parameters found in Table S5. Final $\chi^2$-values shown beneath each model. (C) SAXS profiles with model fits and residuals for each complex colored as in panel (A-B). Scattering curves are offset for clarity.

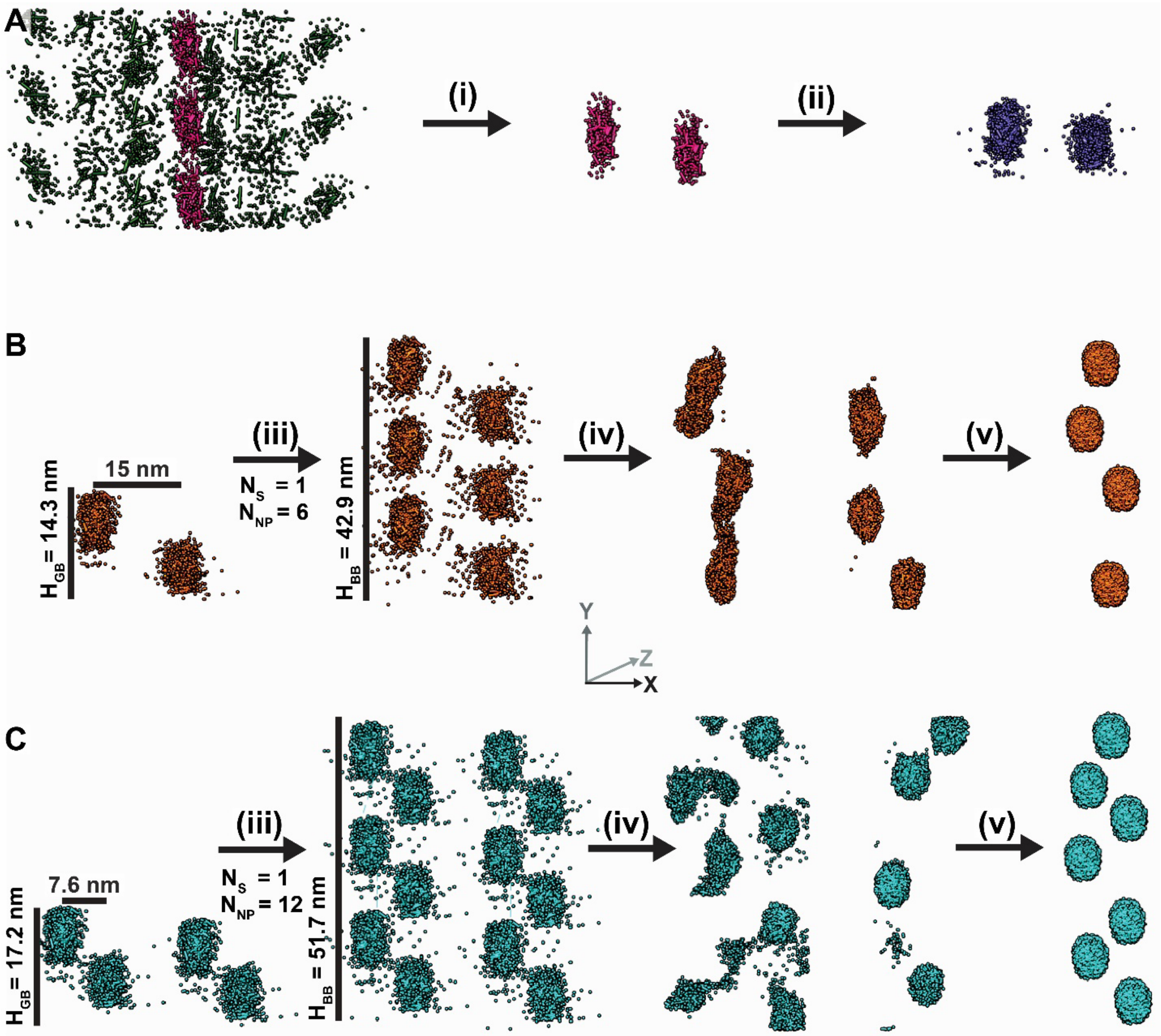


**Figure S16.** Schematic demonstration of adapted SASHEL *ab initio* modeling strategy with labeling of important definitions for stack building block heights ($H_{BB}$), inter-AuNP block heights ($H_{GB}$), number of stacks ($N_S$), and number of AuNPs ($N_{NP}$). The stepwise methodology is as follows: (A) broad movement of 2000 initial dummy atoms from initial guess core-shell model (green) resulting in a densely packed strand of dummy-atom clusters, each representative of an AuNP (pink). (i) One clearly defined cluster is selected and duplicated to give an AuNP pair, (ii) then the pair of AuNPs is expanded to 1000 dummy atoms per AuNP (purple). Each AuNP in the pair is maneuvered into desired initial geometries as determined experimentally, representing one $H_{GB}$ unit, unique for (B) $(GT)_{15}$-AuNP-SWCNTs and (C) $(GT)_6$-AuNP-SWCNTs. (iii) The $H_{GB}$ is replicated in one direction using symmetrical boundary conditions an integer (*k*) number of times to produce a total stack height, $H_{BB} = k*H_{GB}$, using parameters in Table S5. (iv) The starting model is then used to produce ab initio models representing two neighboring ssDNA-AuNP-SWCNTs. (v) The single ssDNA-AuNP-SWCNT showing the clearest AuNP spacing is selected and the regions of electron density are replaced with denser 3000 dummy-atom clusters (AuNPs).

**Table S4.** AuNP starting strand distances series parameters and fitting results for SASHEL modeling.

| | **Starting AuNP Strand Distances (nm)** | 10 | 15 | 20 |
|---|---|---|---|---|
| **$\chi^2$-value of Fit** | $(GT)_{15}$-AuNP-SWCNT | 2.10 | 0.27 | 0.99 |
| | $(GT)_6$-AuNP-SWCNT | 6.72 | 1.19 | 1.35 |

**Table S5.** Stack parameters for SASHEL modeling.

| | **Number of Stacks ($N_S$)** | 16 | 8 | 5 | 4 | 3 | 3 | 2 | 2 | 2 |
|---|---|---|---|---|---|---|---|---|---|---|
| | **AuNPs per Stack ($N_{NP}$)** | 2 | 4 | 6 | 8 | 10 | 12 | 14 | 16 | 18 |
| **Total Stack Height (nm)** | $(GT)_{15}$-AuNP-SWCNT | 14.3 | 28.6 | 42.9 | 57.2 | 71.6 | 85.9 | 100.2 | 114.5 | 128.8 |
| | $(GT)_6$-AuNP-SWCNT | 17.2 | 34.5 | 51.7 | 69.0 | 86.2 | 103.5 | 120.7 | 137.9 | 155.2 |
| **$\chi^2$-value of Fit** | $(GT)_{15}$-AuNP-SWCNT | 284.2 | 14.8 | 3.11 | 0.51 | 0.70 | 3.37 | 0.97 | 0.27 | 0.41 |
| | $(GT)_6$-AuNP-SWCNT | 99.8 | 9.77 | 1.19 | 4.60 | 1.05 | 0.34 | 0.96 | 0.48 | 0.24 |

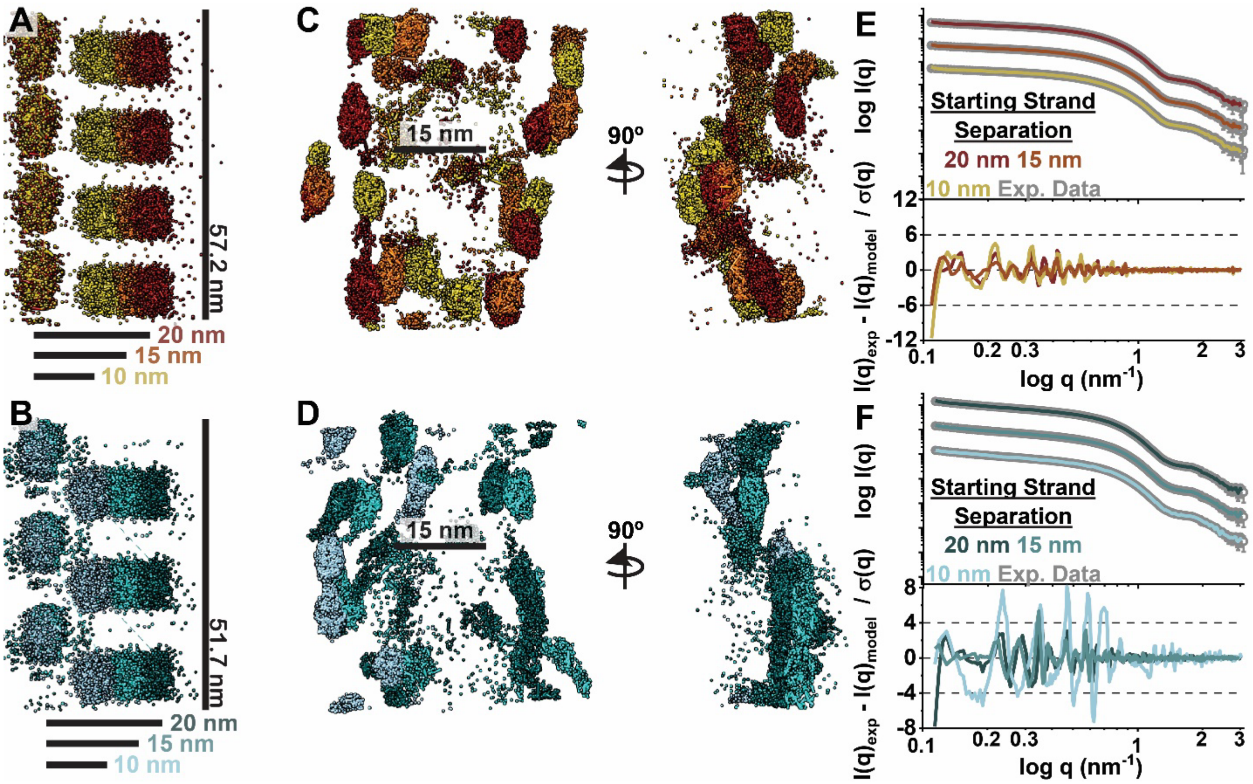


**Figure S17.** *Ab initio* modeling series for different starting AuNP-tagged ssDNA strand distances of 10, 15, and 20 nm via SASHEL, with full parameters shown in Table S4. Starting models for (A) $(GT)_{15}$-AuNP-SWCNTs and (B) $(GT)_6$-AuNP-SWCNTs. *Ab initio* modeling results for (C) $(GT)_{15}$-AuNP-SWCNTs and (D) $(GT)_6$-AuNP-SWCNTs. SAXS profiles with model fits and residuals for (E) $(GT)_{15}$-AuNP-SWCNTs and (F) $(GT)_6$-AuNP-SWCNTs with each complex colored as in panel (A) and (B), respectively. Scattering curves are offset for clarity.

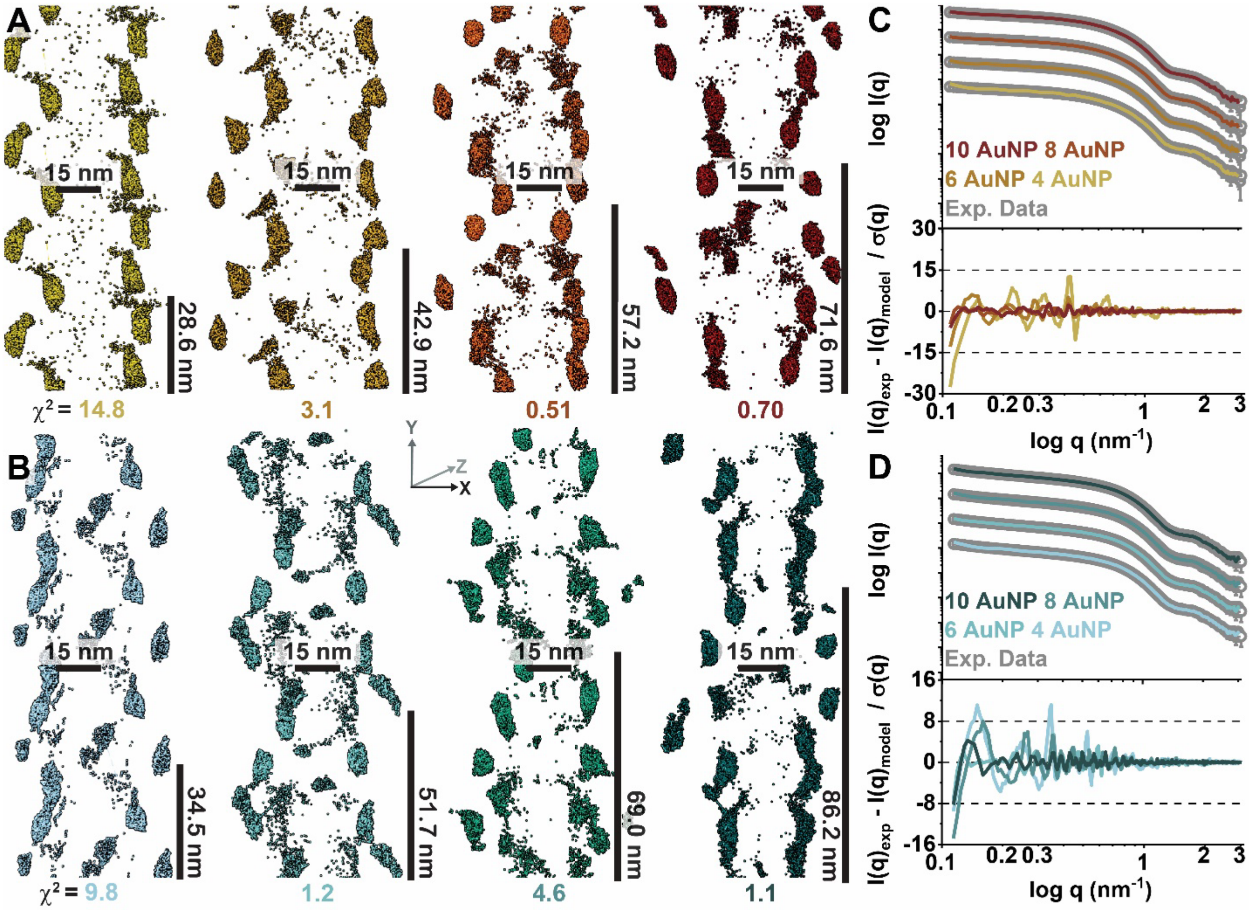


**Figure S18.** *Ab initio* modeling series results for (A) $(GT)_{15}$-AuNP-SWCNTs and (B) $(GT)_6$-AuNP-SWCNTs made via SASHEL for 4 to 10 AuNPs per inter-AuNP block height ($H_{GB}$), as defined in SI Methods and Figure S16. $\chi^2$-values are shown below each model and starting parameters are shown in Table S5. SAXS profiles with model fits and residuals for (C) $(GT)_{15}$-AuNP-SWCNTs and (D) $(GT)_6$-AuNP-SWCNTs, with each sample colored as in panel (A) and (B), respectively. Scattering curves are offset for clarity.

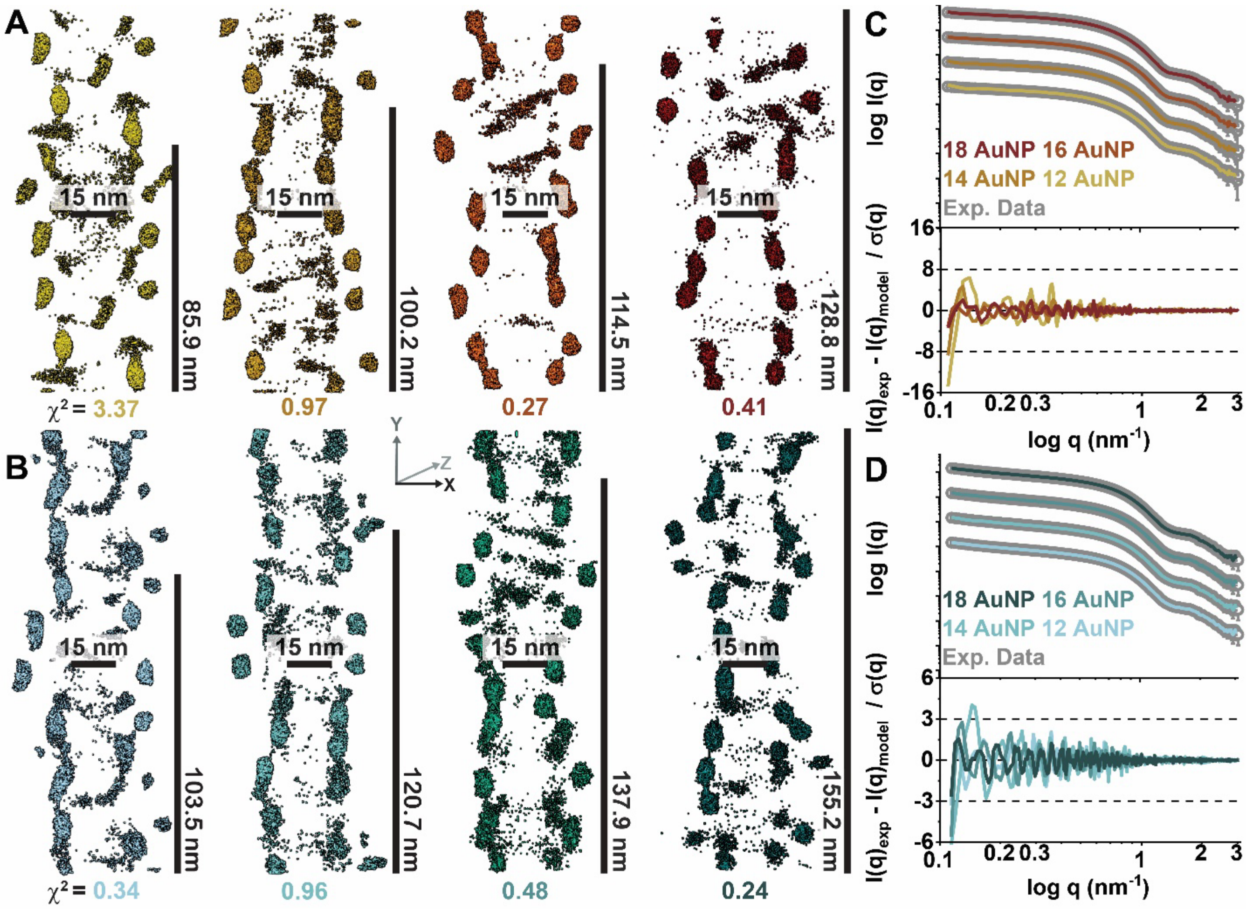


**Figure S19.** *Ab initio* modeling series results for (A) $(GT)_{15}$-AuNP-SWCNTs and (B) $(GT)_6$-AuNP-SWCNTs made via SASHEL for 12 to 18 AuNPs per inter-AuNP block heights ($H_{GB}$), as defined in SI Methods and Figure S16. $\chi^2$-values are shown below each model and starting parameters are shown in Table S5. SAXS profiles with model fits and residuals for (C) $(GT)_{15}$-AuNP-SWCNTs and (D) $(GT)_6$-AuNP-SWCNTs, with each sample colored as in panel (A) and (B), respectively. Scattering curves are offset for clarity.

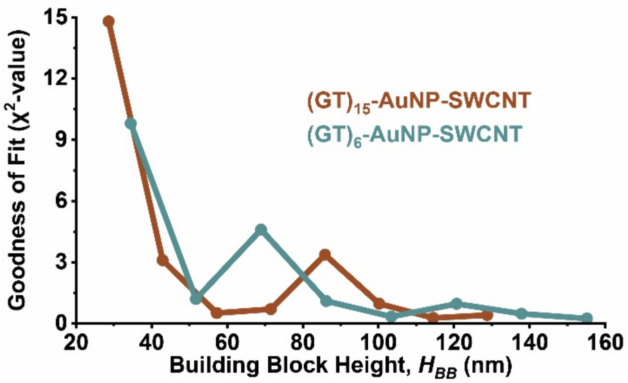


**Figure S20.** *Ab initio* modeling series results for $(GT)_{15}$-AuNP-SWCNTs and $(GT)_6$-AuNP-SWCNTs made via SASHEL showing the goodness of fit ($\chi^2$-values) as a function of stack building block heights ($H_{BB}$). All parameters are shown in Table S5.

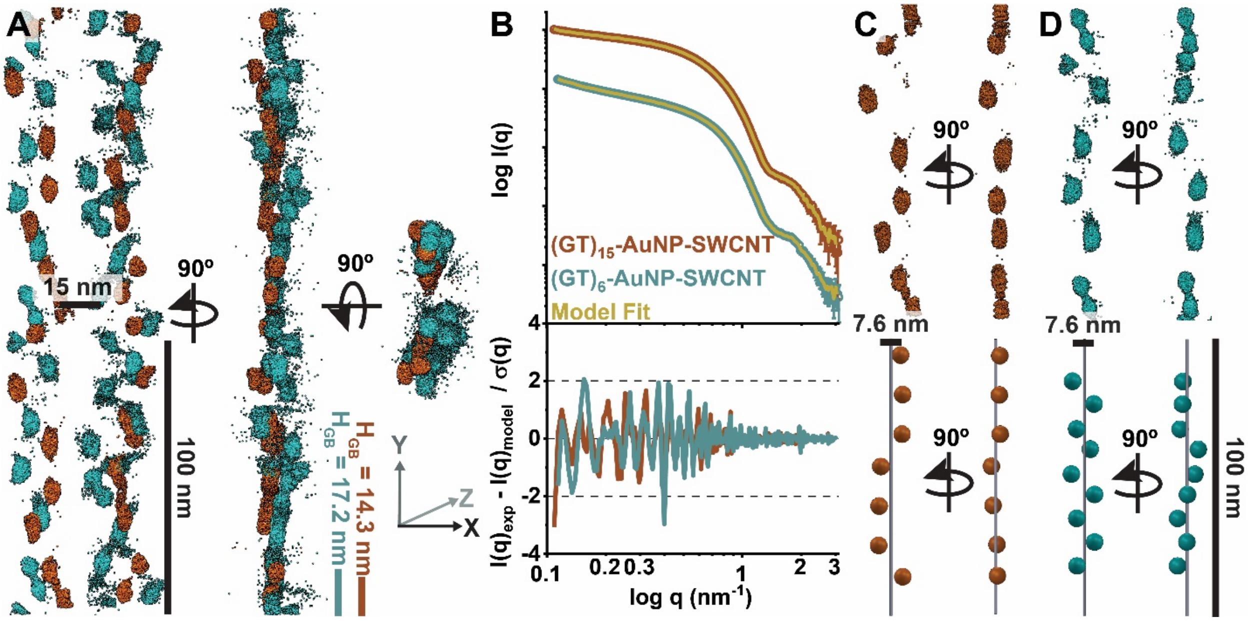


**Figure S21.** *Ab initio* modeling of ssDNA-AuNP-SWCNT complexes reveals periodic ordering of corona phase, using methodologies adapted from the SASHEL software for elongated nanoscale systems (see SI Methods). Both $(GT)_{15}$- and $(GT)_6$-AuNP-SWCNTs were best captured by parallel lines of AuNPs (SWCNTs) with the (A) best-fit models for $(GT)_{15}$-AuNP-SWCNTs (red) and $(GT)_6$-AuNP-SWCNTs (blue) containing 16 or 36 AuNPs per repeating stack building block heights ($H_{BB}$), respectively. Additional parameters for inter-AuNP block heights ($H_{GB}$), number of stacks ($N_S$), and number of AuNPs ($N_{NP}$) used to produce the models can be found in Table S5. Noise reduction of the models was implemented for clarity by removing some un-clustered dummy atoms accounting for 8% of the total, as shown in Figure S23. (B) SAXS profiles with model fits (gold) and residuals for each complex colored as in panel (A). Scattering curves are offset for clarity. (C-D) Comparison of isolated, individual ssDNA-AuNP-SWCNTs from *ab initio* modeling (top) and theoretical 3D diagrams (bottom) produced from distances obtained from *P(r)* functions for (C) $(GT)_{15}$-AuNP-SWCNTs and (D) $(GT)_6$-AuNP-SWCNTs. Diagrams are scaled and colored to match that of panel (A).

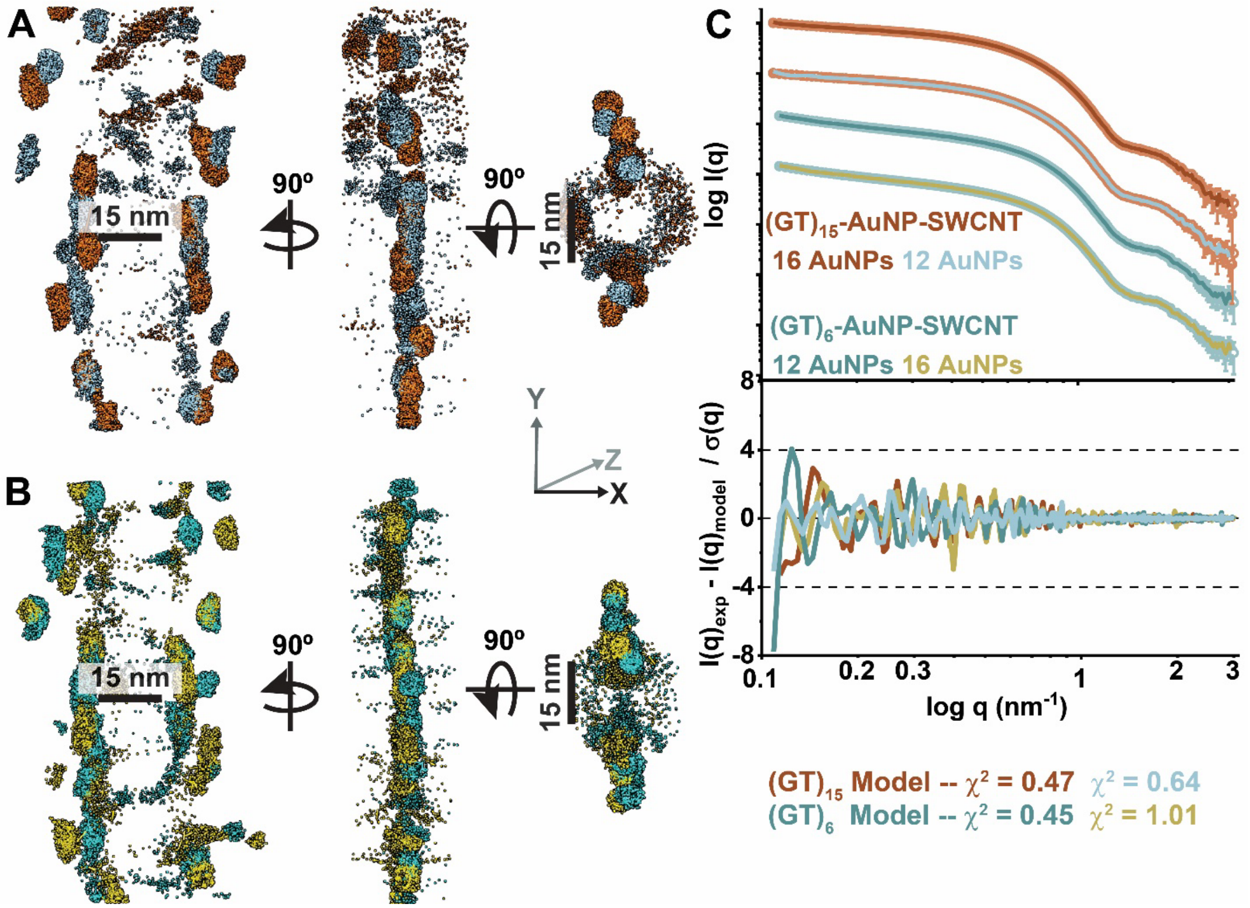


**Figure S22.** *Ab initio* modeling results for switching best-fit models for (A) $(GT)_{15}$-AuNP-SWCNTs and (B) $(GT)_6$-AuNP-SWCNTs. $\chi^2$-values shown for each model. (C) SAXS profiles with model fits and residuals for $(GT)_{15}$- and $(GT)_6$-AuNP-SWCNT complexes with fits colored as in panel colored as in panels (A-B). Scattering curves are offset for clarity.

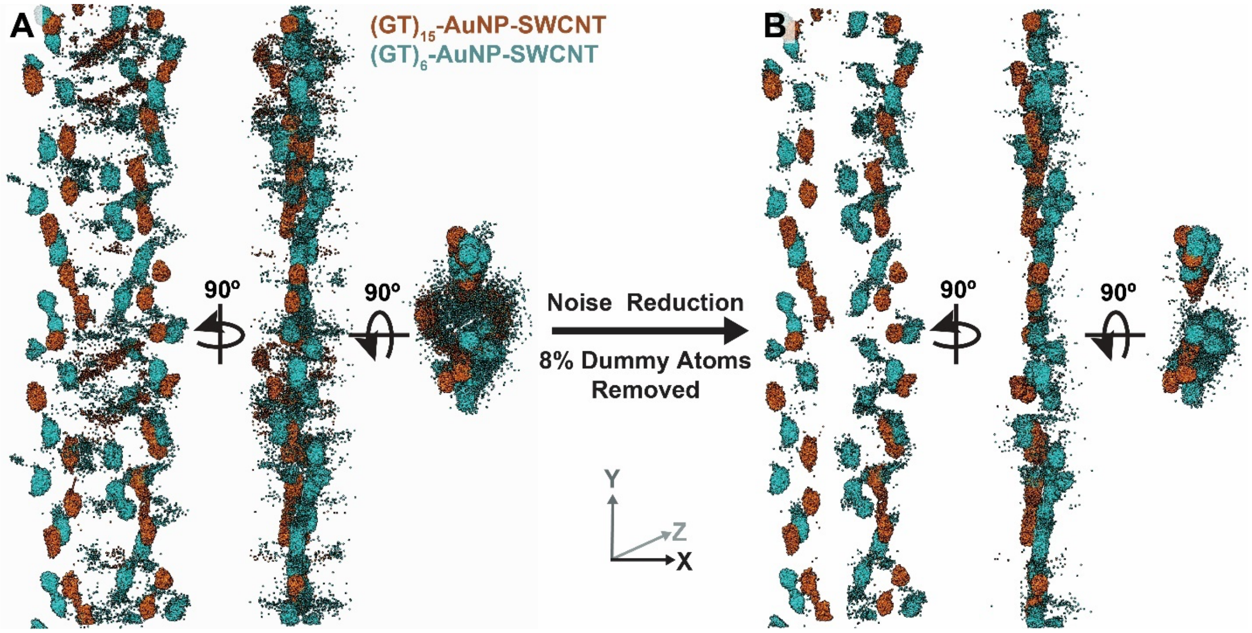


**Figure S23.** Noise reduction of best fit *Ab initio* modeling results by removing some un-clustered dummy atoms accounting for 8% of the total. Models are shown going from (A) original models for $(GT)_{15}$- and $(GT)_6$-AuNP-SWCNT to (B) noise reduced models.
